# Membrane cholesterol regulates inhibition and substrate transport by the glycine transporter, GlyT2

**DOI:** 10.1101/2022.08.31.506132

**Authors:** Zachary J. Frangos, Katie A. Wilson, Heather M. Aitken, Ryan Cantwell Chater, Robert J. Vandenberg, Megan L. O’Mara

## Abstract

Membrane cholesterol binds to and modulates the function of the specific SLC6 transporters. Here we investigate how cholesterol binds to and modulates the rate of glycine transport by the SLC6 glycine transporter GlyT2, and how this impacts lipid inhibition of GlyT2. Bioactive lipid inhibitors of GlyT2 are analgesics that bind to the lipid allosteric site of the outward facing GlyT2 conformation that is accessible from the extracellular solution. Using molecular dynamics simulations, mutagenesis and cholesterol depletion experiments, we show that bioactive lipid inhibition of glycine transport is modulated by the recruitment of membrane cholesterol to a cholesterol binding site formed by transmembrane helices 1, 5 and 7. Recruitment involves cholesterol flipping from its membrane orientation, and insertion of the 3’ hydroxyl group into the cholesterol binding cavity to interact with the base of the lipid allosteric site and the bound inhibitor. The recruitment of membrane cholesterol by allosteric GlyT2 inhibitors is a potential avenue for the development of high-potency, specific pain analgesics and could provide alternative therapeutics that target GlyT2 and other SLC6 neurotransmitter transporters.

## Introduction

The glycine transporter 2 (GlyT2) is a member of the SLC6 family of neurotransmitter transporters that plays an important role in regulating glycine concentrations in inhibitory synapses of the ascending pain pathway. After presynaptic release of glycine, GlyT2 transports glycine from the synaptic cleft and back into the pre-synaptic neuron where it can be recycled into synaptic vesicles to maintain glycinergic neurotransmission. SLC6 transporters are secondary active transporters that exploit Na^+^ gradients to drive the transport of substrate across cell membranes through a conserved general mechanism (*1*). Substrate and ions bind to an outward-open conformation of the transporter. The transporter then undergoes a series of conformational changes resulting in an inward-open conformation that allows the release of substrate to the intracellular environment.

GlyT2 and other SLC6 transporters localise in cholesterol-enriched lipid raft domains in the plasma membrane (*2*, *3*) and their activity is modulated by membrane cholesterol concentration (*4*). Recent crystal and cryo-EM structures, as well as molecular dynamics (MD) simulations, have identified five cholesterol binding sites (CHOL1-5) in other SLC6 transporters including the drosophila dopamine transporter (dDAT), human dopamine transporter (hDAT) and human serotonin transporter (SERT) (*5*–*9*).Depletion of membrane cholesterol by methyl-β-cyclodextrin (mβCD) reduces transporter activity (*4*, *10*–*14*), while experimental and MD work suggests that the presence of cholesterol changes the transporter conformational equilibrium, favouring the outward facing conformation (*7*, *15*–*17*). Further coarse-grained MD studies comparing the lipid annulus of select SLC6 transporters show transporter-specific differences in cholesterol occupancies across these crystallographic binding sites (*18*).

In dDAT, CHOL1 is formed by TMs 1a, 5 and 7 which also form part of the transporter’s core domain that undergoes significant conformational changes during the transition from the outward-facing to inward-facing state of the transporter (*7*). Thus, cholesterol bound to the CHOL1 site is mostly likely to be responsible for the regulatory actions of cholesterol on transporter function. This is supported by MD simulations of hDAT, where cholesterol bound in CHOL1 prevents conformational changes that mediate intracellular gating such as unwinding of TM5, an essential component in the early stages of the transition to the inward-facing state (*19*, *20*). Furthermore, treatment with mβCD and introduction of polar residues into CHOL1 of hSERT through mutagenesis disrupts cholesterol binding and results in a favouring of the inward-facing conformation (*15*, *17*). This has led to the notion that cholesterol bound in CHOL1 stabilises the outward-facing state of neurotransmitter sodium symporters (NSS) and forms a dynamic part of the transport cycle, regulating transitions between the outward and inward conformations.

Membrane cholesterol content has also been shown to alter the affinity of NSS ligands, based on the conformational state they preferentially bind. Depleting membrane cholesterol shifts the conformational equilibrium of transporters towards the inward-facing state and enhances the activity of compounds that bind to this conformation and decreases the activity of ligands that bind the outward-facing conformation (*4*, *17*). Similarly, supplementing the membrane with cholesterol shifts the conformational equilibrium towards the outward-facing state and increases the activity of compounds that preferentially bind this conformation (*16*).

Inhibitors of GlyT2 slow the removal of glycine from the synaptic cleft, prolonging glycinergic transmission to inhibit pain (*2*, *3*). We have previously shown that N-*ocyl* amino acids are a potent class of lipid-based GlyT2 inhibitors (*21*, *22*). These inhibitors bind to a site (referred to as the lipid allosteric site; LAS) that is accessible from the extracellular solution in the outward facing conformation (*23*). In subsequent work we have shown that the activity of these lipid inhibitors is influenced by the depth of penetration of the lipid tail into the LAS (*24*). Nevertheless, the interplay between lipid inhibitor and cholesterol binding to GlyT2 is unknown. In the present work we use a combination of molecular dynamics simulations, cholesterol depletion techniques and mutagenesis to examine the interactions between bioactive lipid inhibitors bound to GlyT2 and how membrane cholesterol influences these interactions. We demonstrate that glycine transport by GlyT2 and its sensitivity to bioactive lipid inhibition is influenced by interactions with cholesterol molecules recruited from the membrane to a specific site on GlyT2.

## Results

### The LAS is only accessible in the outward facing conformation of GlyT2

To better understand if lipid inhibitor binding is specific to a single conformation of GlyT2, we compared the structure of the LAS in the inward and outward facing conformations of GlyT2. We used our previously characterized outward facing conformation (23, 24, 39) and developed an homology model of inward-facing GlyT2, using the inward-facing cryoEM structure of human SERT as a structural template (PDB ID: 6DZZ) (*25*). Triplicate 500 ns united atom simulations were performed on both the inward and outward facing GlyT2 models and the LAS pocket volume was calculated for both conformations. Consistent with previous studies (*23*, *24*), throughout all replicate simulations of outward facing GlyT2 simulations, the LAS is accessible from the extracellular solution. The volume of the LAS cavity is between 750–1750 Å^3^ for 60% of the simulation, and the minimum volume always exceeds 500 Å^3^ (Figure 1A). In contrast, the LAS volume in the inward conformation is substantially smaller. For 77% of the total 1500 ns, the LAS volume <750 Å^3^ (Figure 1B). This includes an extended period (510 ns, or 34% of the total simulation time) in which the LAS is completely absent. Notably, this reduced volume LAS is not accessible from the extracellular solution. This indicates that a functional LAS capable of inhibitor binding is only formed in the GlyT2 outward facing conformation.

**Figure 1.**
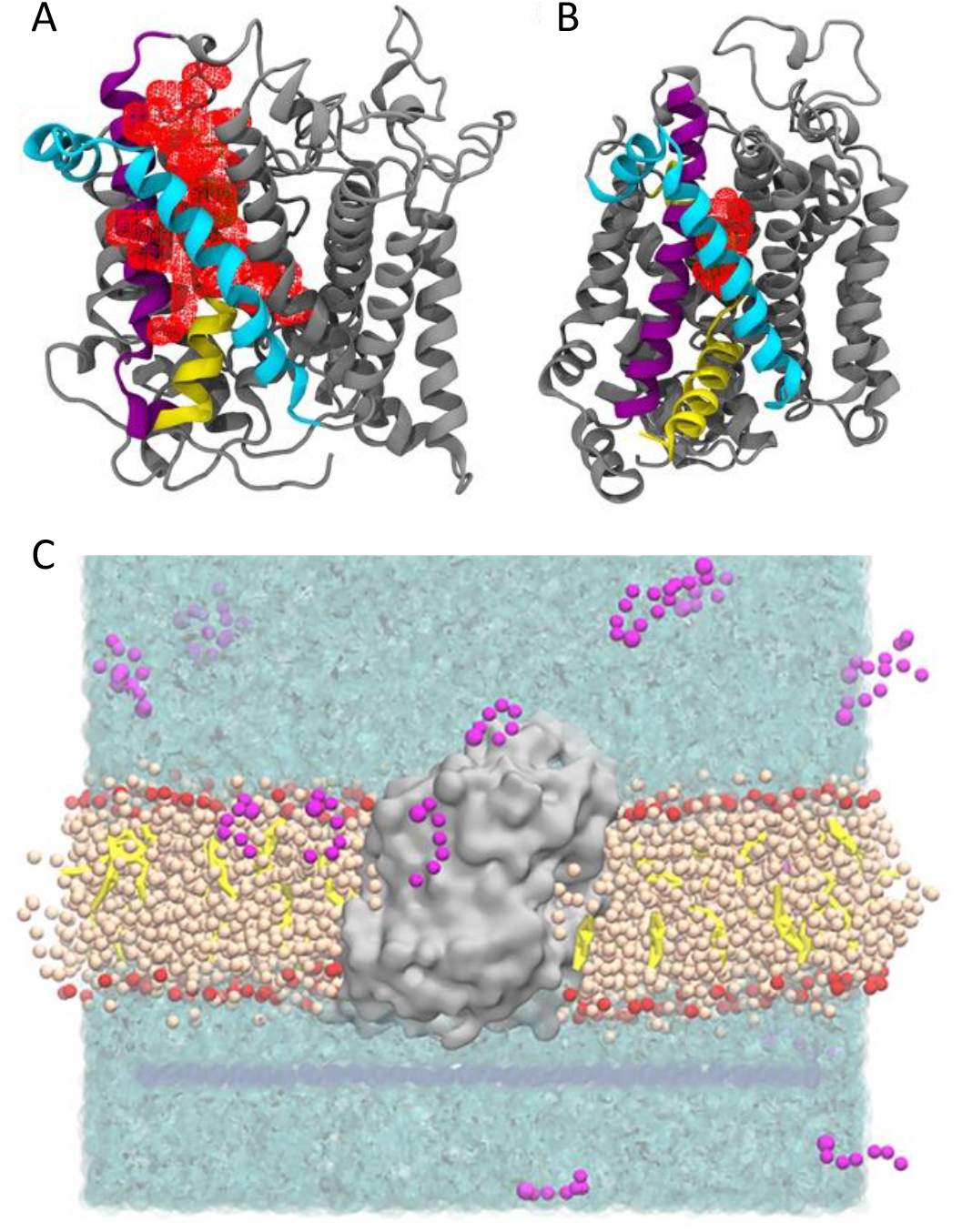
Homology models of GlyT2 in the outward and inward conformations. Pocket volume (red mesh) for the LAS in the A) outward open and B) inward open conformations of GlyT2. Important transmembrane domains are shown including TM1 (yellow), TM5 (cyan), and TM7 (purple). **C)** Coarse grained molecular dynamics simulation setup showing GlyT2 (gray) embedded in an 80% POPC (tan) 20% cholesterol (yellow) membrane, with 20 molecules of a given species of lipid inhibitor (magenta) in the aqueous solution (cyan).

### Lipid inhibitors interact with membrane cholesterol

To investigate the spontaneous binding of lipid inhibitors to the GlyT2 LAS, coarse-grained (CG) MD simulations were performed on the GlyT2 outward facing conformation embedded in an 80% POPC 20% cholesterol membrane, with 20 molecules of a given species of lipid inhibitor in the aqueous solution, as shown in Figure 1C. Four different lipid inhibitors were examined in these spontaneous binding simulations, namely oleoyl L-tryptophan (OLTrp), oleoyl L-serine (OLSer), oleoyl L-leucine (OLLeu) and oleoyl L-lysine (OLLys). The majority of the lipid inhibitors adsorbed to the surface of the membrane or formed micelles before adsorbing and partitioning into the membrane (Figure EVA1). The partitioning of the lipids into the membrane did not significantly alter overall membrane properties (28) (Table EV1). In one replicate simulation, a single OLLeu lipid inhibitor entered the LAS from the extracellular solution after 0.05 μs of simulation, where it remained bound for a further 9.3 μs, before dissociating from the LAS and entering the membrane. While in the LAS, OLLeu interacted with residues from EL4, TM7, and TM8 (Figure 2A and Table EV2). In particular, OLLeu interacts with F567, I524, M527 and W563 for >8 μs and with W215 and Y550 for >6 μs during the 10 μs simulation. The combined effect of these interactions is likely responsible for stabilising OLLeu in this site. Notably, Y550 and W563 have been previously shown through mutagenesis experiments to affect the activity of lipid inhibitors on GlyT2 (*23*). When considered together, this demonstrates that a lipid inhibitor can spontaneously bind from solution to the GlyT2 LAS, and that the binding pose adopted is consistent with that observed in previous docking and atomistic simulations (*23*, *24*).

**Figure 2.**
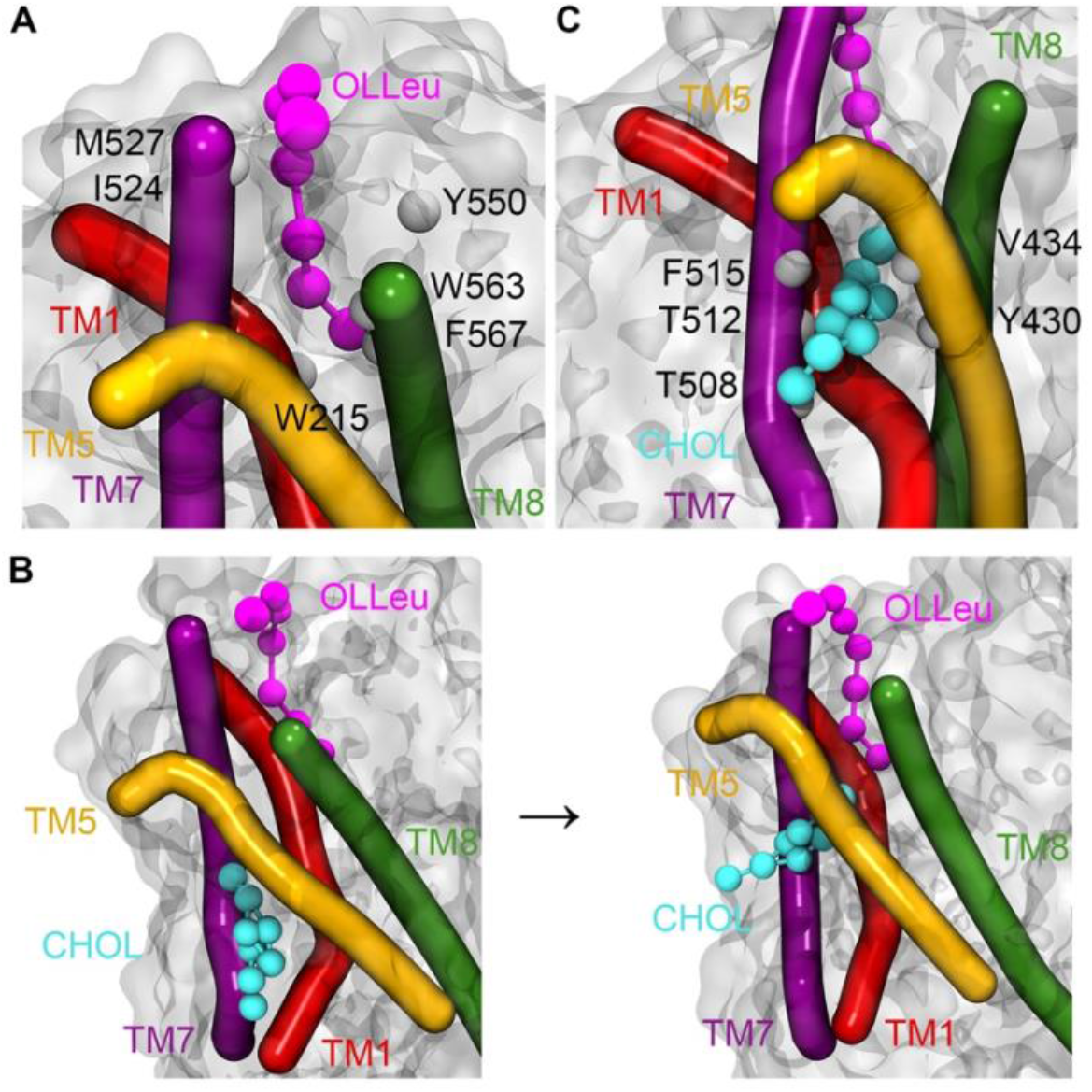
Representative structure showing OLLeu and associated CHOL bound in the LAS of GlyT2 from the CG spontaneous binding simulations. A) OLLeu bound in the LAS with residues that are in contact with the lipid inhibitor for >60% of the total simulation time highlighted. B) CHOL initial bound in the dDAT CHOL1 site before flipping orientations to burrow between TMs 5 and 7 at the bottom of the LAS. C) CHOL bound at the bottom of the LAS site with residues that are in contact with the lipid inhibitor for >75% of the simulation time that CHOL is interacting with GlyT2 highlighted. Important transmembrane domains are shown including TM1 (red), TM5 (yellow), TM7 (purple) and TM8 (green) with cholesterol (cyan) and OLLeu (magenta) presented as spheres.

Whilst bound in the LAS, a previously unseen interaction occurs between OLLeu and membrane cholesterol. After 1 μs of CG simulation, a molecule of cholesterol within the membrane associates with TMs 1a, 5 and 7 in a similar orientation to that observed for cholesterol bound at the CHOL1 site of dDAT. However, after 2.6 μs, the cholesterol molecule flips to adopt the opposite orientation in the membrane, burying deeper into a cavity between TMs 1, 5 and 7 of GlyT2, such that the 3’ hydroxyl group of cholesterol forms a close association with the terminal end of the bioactive lipid tail (Figure 2B) and the isooctyl tail is partitioned into the phospholipid acyl tails and oriented at a near-perpendicular angle to the plane of the membrane. Cholesterol remains bound to the cavity for 5 μs of CG simulation. In this flipped orientation, the cholesterol molecule is in direct contact with the bound OLLeu for 30% of the time cholesterol is bound. On entering the cholesterol binding cavity formed by TMs 1, 5, and 7, the cholesterol molecule is within 6.0 Å of Y430, L433, V434, T508, A511, T512 and F515 for >75% of the time it is bound (Figure 2C and Table EV3). The combined effect of these interactions is likely responsible for stabilising cholesterol in this site. After 5 μs of CG simulation, cholesterol dissociates from the binding cavity and reorients within the bulk of the membrane. Without cholesterol bound between TMs 1a, 5 and 7, OLLeu dissociates, suggesting that the interaction between OLLeu and membrane cholesterol may be an important factor in the binding and mechanism of inhibition of these lipids.

To further understand how the function of GlyT2 is modulated by the binding of the lipid inhibitor to the extracellular allosteric pocket, a representative frame from the CG simulation containing both bound cholesterol and OLLeu was backmapped to atomistic detail, and the system simulated in triplicate for a further 500 ns using the GROMOS 54a7 forcefield (*26*). In all three replicates, the lipid inhibitor remained in the LAS located between EL4, TM5, TM7 and TM8 (Figure 3). While there were variations in the precise orientation of the lipid inhibitor tail across the three replicate simulations, the lipid inhibitor formed direct contacts (within 4 Å) with P218, L437, I520, V523 M527, F562, W563, I566, F567 and M570 for >50% of the combined 1500 ns of the backmapped atomistic simulations (Table EV4). This is consistent with the contact residues observed in the CG simulations and those previously proposed in a combined mutagenesis/docking study (*23*).

**Figure 3.**
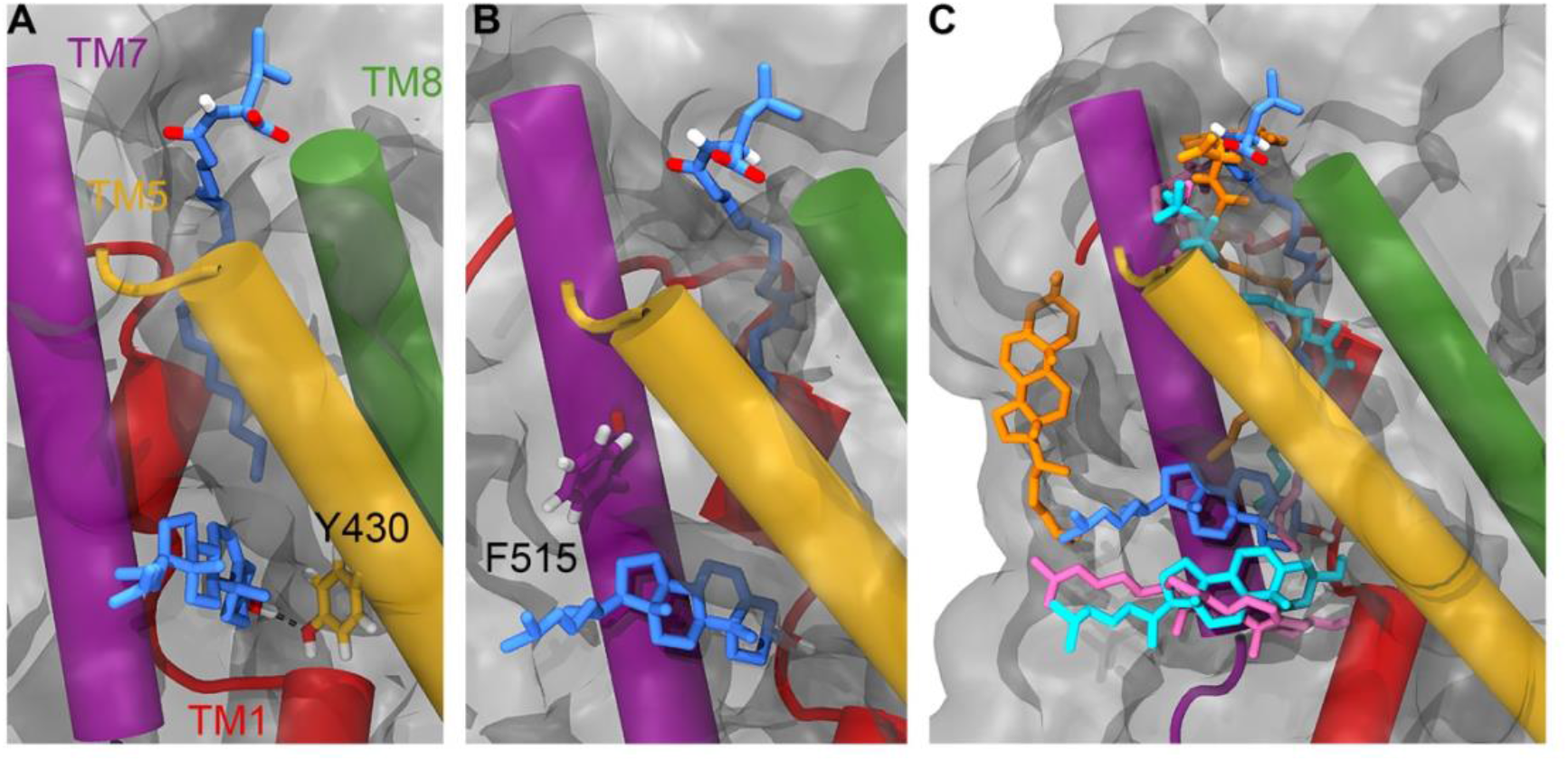
Position of the lipid inhibitor and associated CHOL at the end of 500 ns atomistic simulations. **A)** Position of Y430 (yellow liquorice) relative to OLLeu (blue liquorice, upper) and bound cholesterol (blue liquorice, lower). **B)** Position of F515 (purple liquorice) relative to OLLeu (blue liquorice, upper) and bound cholesterol (blue liquorice, lower). **C)** Overlay of the relative position of OLLeu (blue), OLLys (pink), OLCarn (cyan) and OLTrp (orange) and cholesterol in the LAS from 500 ns simulations. Cholesterol is shown in the same colour as the lipid inhibitor from the corresponding simulation. Important transmembrane domains are shown including TM1 (red), TM5 (yellow), TM7 (purple) and TM8 (green).

The atomistic simulations provide a deeper biochemical insight into how the bound cholesterol is interacting with GlyT2 and the lipid inhibitor. While cholesterol remains bound in the LAS for ~1050 ns of the combined 1500 ns of simulation time, the precise interactions formed by the bound cholesterol molecule are dependent on the relative orientation of the OLLeu lipid inhibitor in each of the three replicate simulations. When the inhibitor tail is extended, the side chain of Y430, located at the base of the extracellular allosteric pocket, is oriented into the cavity between TMs 1a, 5 and 7 (Figure 3). In this orientation, cholesterol binding is primarily stabilised through its interaction with Y430, through both hydrogen bonding (with the 3’ hydroxyl) and π-stacking interactions. Without this reorientation of Y430 cholesterol dissociates from the LAS. Interactions between Y430 and cholesterol occur for 65% (975 ns) of the combined 1500 ns simulation. Cholesterol binding is further stabilised through a dynamic hydrogen bonding network with residues within the TMs 1a, 5, and 7 cavity, mediated by V205 and through transient contacts with the nearby residues T427 and T573. Residues C507, T508, A511, T212 and F515 are also in close proximity to bound cholesterol molecule (Table EV5). Overall, these atomistic simulations suggest that the position of the lipid tail and reorientation of Y430 (Figure 3A) dictate the binding of cholesterol.

As the spontaneous binding of a single lipid inhibitor to the LAS of GlyT2 is a statistically rare event in a simulation system containing 20 inhibitors, the interactions between cholesterol and three different lipid inhibitors, OLLys, oleoyl-L-carnitine (OLCarn), and OLTrp, were investigated by docking the inhibitors to the LAS in the backmapped GlyT2 system with bound cholesterol. Consistent with the observations for OLLeu system, in the presence of OLLys or OLCarn, cholesterol remains near the terminal end of the lipid inhibitor tail, in the cavity between TMs 1a, 5 and 7 for >1000 ns of the combined atomistic simulation time for each lipid inhibitor (Figure 3 and Table EV4). It is noteworthy that the bound conformation of cholesterol depends on the species of inhibitor bound at the LAS. Similar to OLLeu, when OLCarn or OLLys is bound to the LAS, the cholesterol molecule lies nearly perpendicular to the membrane and interacts with V205, Y430 and F515 for > 50% of the total simulation time (Table EV5). In this position the cholesterol molecule forms hydrogen bonding interactions with V205, Y399, and T481. In contrast, with OLTrp bound at the LAS, cholesterol does not maintain this perpendicular orientation and is bound for less than half of the total simulation time. Here, cholesterol either rapidly dissociates from the cavity and associates with the surface of TM7, or loosely associates with the base of the LAS (Figure 3c). The most notable difference between the binding of the cholesterol-interacting bioactive lipids and OLTrp is the orientation of Y430 (TM5). The tails of OLLys, OLLeu, and OLCarn penetrate deeper into the binding cavity. This deeper inhibitor penetration is associated with the reorientation of the Y430 side chain into the cavity between TMs 1a, 5 and 7. Together, these facilitate the cholesterol hydrogen bonding and stacking interactions that stabilise its binding. In contrast, the tail of OLTrp does not penetrate as deep into the binding site and the side chain orientation of Y430 is unperturbed, preventing formation of the cholesterol binding site. Previous studies have shown the deep penetration of bioactive lipids into the LAS is critical for potent inhibition of GlyT2 (*24*).

### Cholesterol depletion alters the transport activity of GlyT2 and the potency and reversibility of some GlyT2 inhibitors

To explore if the interaction between membrane cholesterol and bioactive lipid inhibitors observed in simulations produces a functional effect, the activity of these compounds was assayed in cholesterol depleted *Xenopus laevis* oocytes. Treatment with MβCD significantly altered the transport activity of GlyT2 with maximal relative transport velocity (V_max_) reduced from 1.00 to 0.59 (p < 0.001; Figure 4 and Table EV6). A trend towards higher apparent glycine affinity was also observed, however this did not reach statistical significance (p = 0.064). These alterations in glycine transport align with previous examinations of the effect of MβCD induced cholesterol depletion on the transport activity of GlyT2, as well as DAT and SERT (*4*, *11*, *14*), indicating efficient cholesterol depletion.

**Figure 4.**
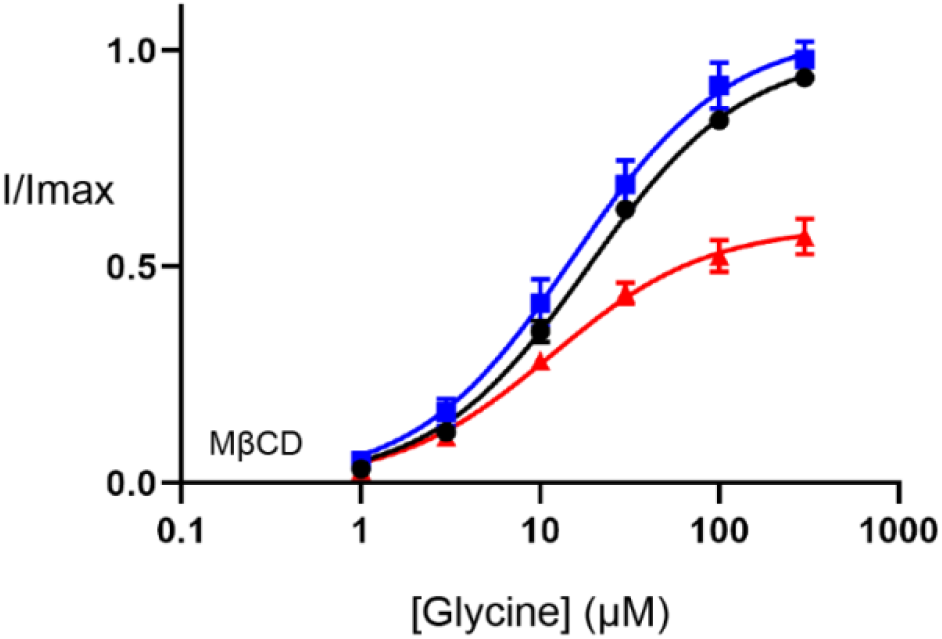
Membrane cholesterol depletion alters the functionality of WT GlyT2 expressed in *Xenopus laevis* oocytes. Transport currents were measured in the presence of increasing glycine concentrations (1-3000 μM) before, and after, treatment with methyl-β-cyclodextrin (MβCD). Baseline responses (black) were performed before incubating oocytes with 0 mM (blue) or 15 mM (red) cyclodextrin for 30 minutes at 32°C. After incubation, oocytes were washed for 10 minutes in recording buffer and glycine concentration-responses repeated. Currents were fit to the modified Michaelis-Menten equation (see methods) and normalised to the V_max_ of baseline responses. Data points are presented as mean ± SEM (n ≥ 5).

Inhibitor concentration-responses were measured to examine the effect of cholesterol depletion on the activity of selected bioactive lipids. Cholesterol depletion does not alter the potency of OLCarn or OLTrp (Figure 5 and Table EV7), but does significantly reduce the potency of OLLys and OLLeu by 2.3 (p = 0.019) and 2.4-fold (p = 0.008), respectively (Figure 5 and Table EV7). No differences were observed in the amount of inhibition produced by application of 3 μM of any of the inhibitors (Table EV7). This data is consistent with MD simulations identifying interactions between cholesterol and the bound OLLys or OLLeu, and the lack of interaction between cholesterol and bound OLTrp. Taken together, these results suggest that the presence of cholesterol enhances the potency of bioactive lipids in a headgroup dependent manner.

**Figure 5.**
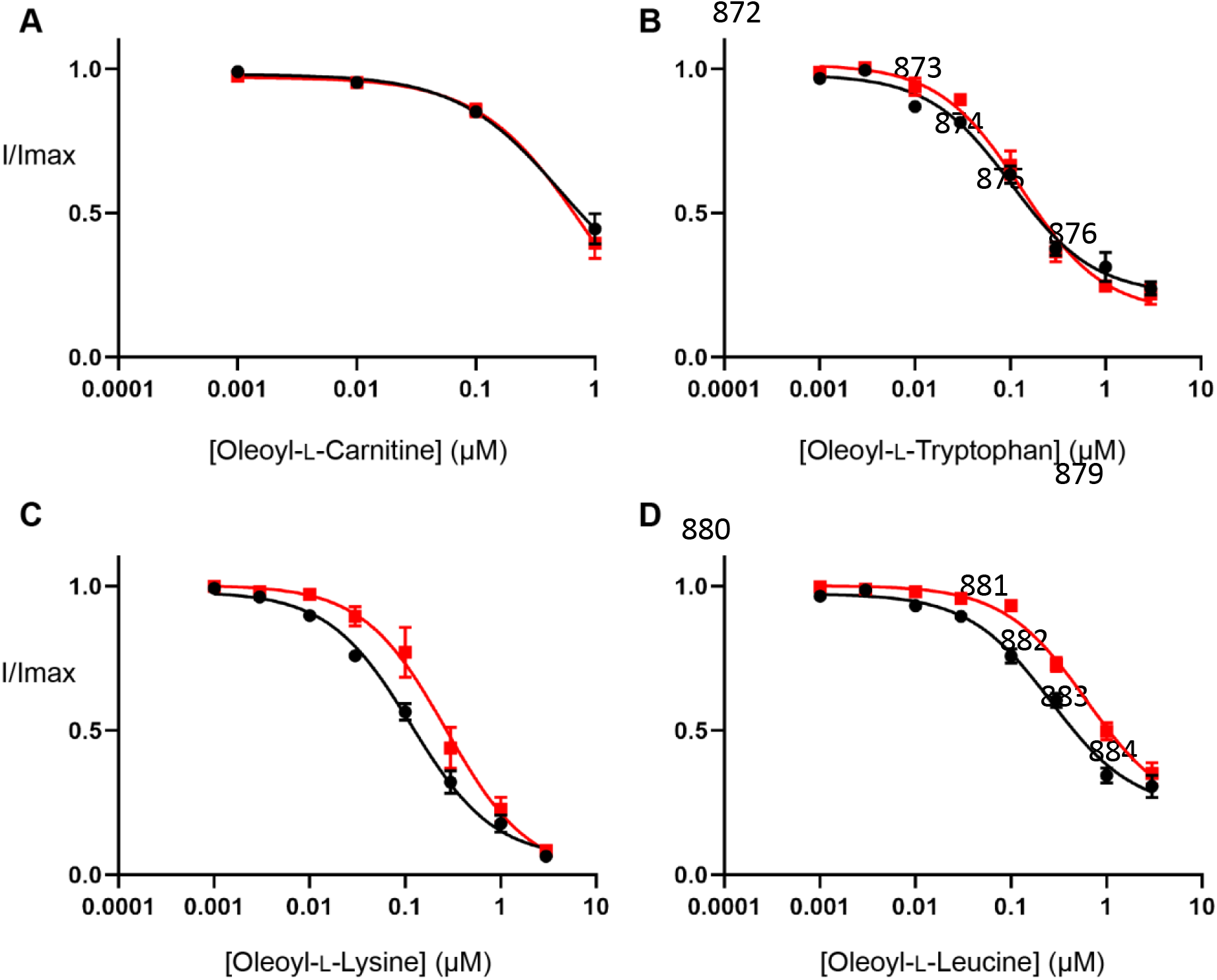
Cholesterol depletion reduces the potency of some bioactive lipids at WT GlyT2 expressed in *Xenopus laevis* oocyte. An EC_50_ concentration of glycine was co-applied with increasing concentrations of (**A**) oleoyl-L-carnitine, (**B**) oleoyl-L-tryptophan, (**C**) oleoyl-L-lysine and (**D**) oleoyl-L-leucine ranging from 1 nM to 3 μM. Concentration-responses performed on control (black) and cholesterol depleted (red) oocytes are shown. Cholesterol depletion was performed by incubating oocytes in 15 mM MβCD for 30 minutes at 32 °C. Raw currents were normalised to currents generated by application of the glycine EC_50_ alone. Data points are presented as mean ± SEM (n ≥ 5).

To examine the influence of membrane cholesterol on the reversibility of GlyT2 inhibition, 30-minute inhibitor washout assays were performed using the IC_50_ of the inhibitor with and without cholesterol depletion. The rate of washout of OLTrp, which is not predicted to interact with cholesterol, was unchanged, and was not reversed (Figure 6 and Table EV8). The transport current recovery half-life following OLLeu application was unchanged, but there was a significant increase in the level of transport current recovery after 30 minutes from 49.4 to 90.7% (p < 0.0001) following cholesterol depletion. Transport current recovery half-lives were unable to be determined for OLCarn and OLLys since recovery from these compounds did not plateau within the timeframe of the assay. Therefore, the level of transport current recovery after 30 minutes was used as a measure of reversibility. Cholesterol depletion significantly increased the transport current recovery 30 minutes after OLCarn treatment from 45.7 to 92.4% (p < 0.0001) (Figure 6 and Table EV8). The similarity in shape of the recovery curves following OLCarn inhibition indicates both conditions are likely to reach the same maximal recovery with sufficient time. Thus, a greater level of transport current recovery at the same timepoint suggests cholesterol depletion reduces the recovery half-life, allowing OLCarn to washout quicker. In contrast, no difference was observed in the shape of the curves, or transport current recovery after 30 minutes, following OLLys application, indicating cholesterol depletion does not alter its reversibility (Figure 6 and Table EV8). Together this data suggests that interactions with membrane cholesterol influences the reversibility of some, but not all, bioactive lipids.

**Figure 6.**
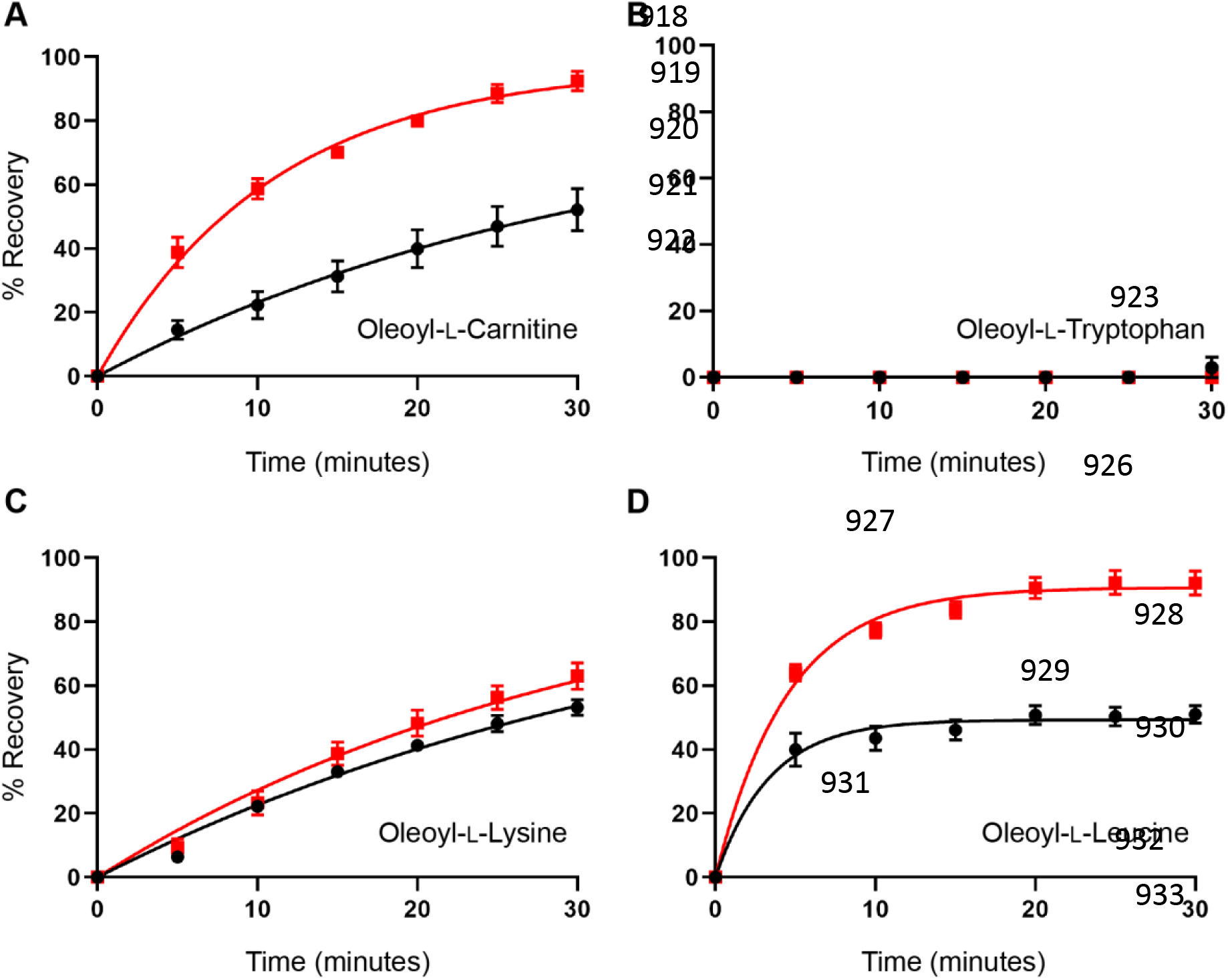
Cholesterol depletion enhances the reversibility of inhibition of WT GlyT2 expressed in *Xenopus laevis* oocytes by some bioactive lipids. Washout time course of (**A**) Oleyol-L-Carn, (**B**) Oleyol-L-tryptophan, (**C**) Oleyol-L-lysine, and (**D**) Oleyol-L-leucine. Membrane cholesterol was depleted by treating oocytes with 15 mM MβCD at 32°C for 30 minutes. Following a 10-minute wash period in recording buffer, an EC_50_ concentration of glycine was co-applied with an IC_50_ concentration of inhibitor for 4 minutes. Following exposure to inhibitors, the EC50 of glycine was re-applied at 5-minute intervals for 30-minutes. % Recovery responses are shown for control (black) and MβCD treated (red) oocytes. Raw currents were normalised to the currents generated by application of the glycine EC_50_ alone. Data points are presented as mean ± SEM (n ≥ 5).

### Mutagenesis of the CHOL1 site mimics cholesterol depletion effects

To further investigate the importance of the CHOL1 site in mediating interactions between GlyT2 inhibitors and membrane cholesterol, site-directed mutagenesis was performed on residues that maintained the longest interactions throughout MD simulations, namely Y430 (Y430F/L), T508 (T508I), T512 (T512A) and F515 (F515V/W) (Figure 7). To examine the importance of cholesterol orientation, L198 (L198A) and I201 (I201V/N) were also mutated. The impact of these mutations were investigated by analysing the effect on inhibition of glycine transport and through MD simulations.

**Figure 7.**
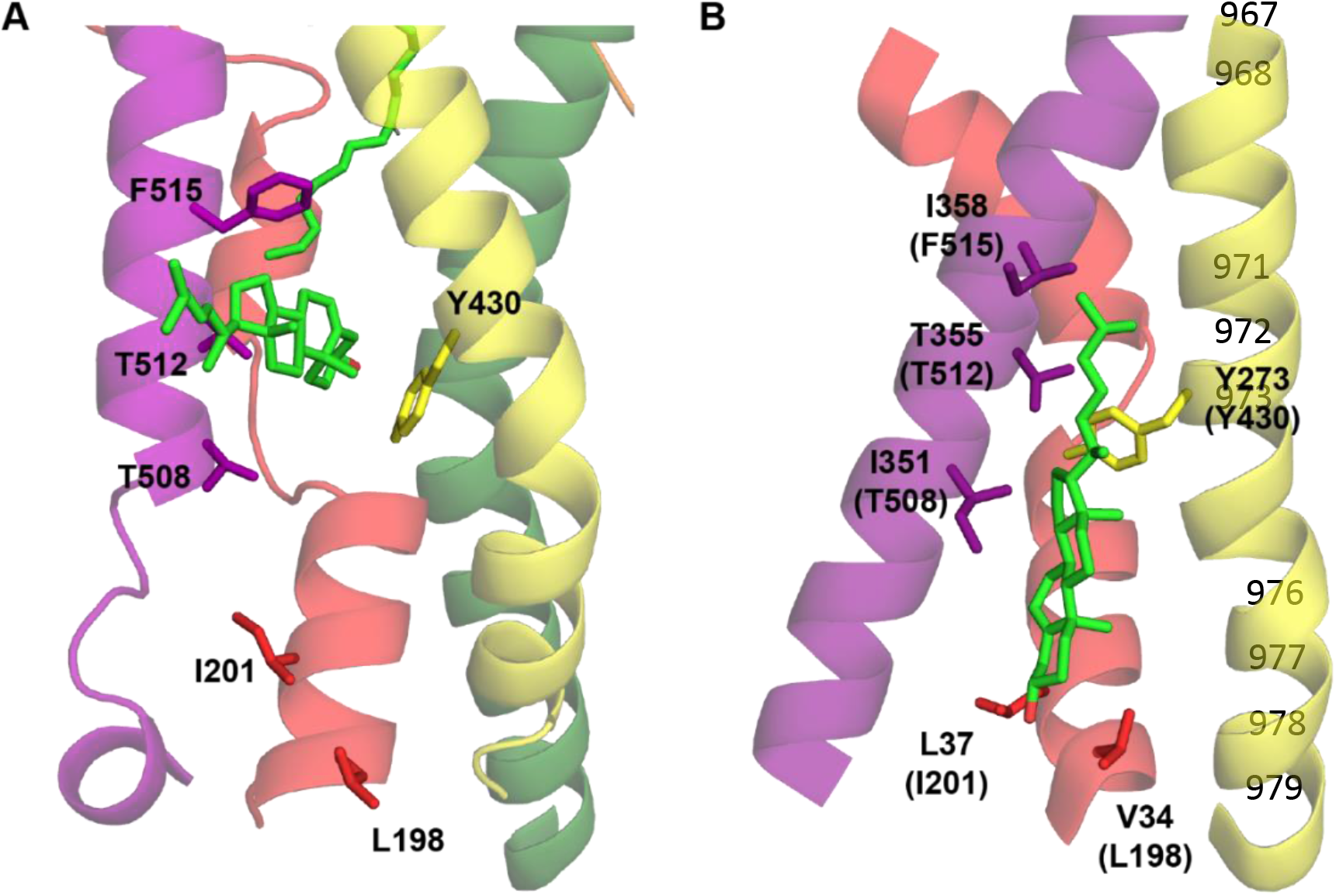
Cholesterol coordinating residues investigated by site-directed mutagenesis. Key stabilising residues of cholesterol in the (**A**) flipped orientation observed in MD simulations and (**B**) non-flipped orientation observed in the dDAT structure (PDB: 4M48) are represented as sticks. GlyT2 and dDAT numbering are used in (**A**) and (**B**) respectively with corresponding GlyT2 residues shown in brackets in (**B**) for clarity. Important transmembrane domains are shown including TM1 (red), TM5 (yellow), TM7 (purple) and TM8 (green) with cholesterol shown as green sticks with oxygen atoms in red. Note the proximity of the cholesterol hydroxyl group to the coordinating residues in each orientation.

All GlyT2 mutants, except I201V/N and T508I, produced reliable and robust glycine-dependent transport currents (Figure 8 and Table EV9). When compared to WT GlyT2, L198A increased the glycine EC_50_ 1.5-fold, from 19 to 28 μM (p = 0.0097). This is consistent with reduced serotonin affinity at the equivalent hSERT mutant (V86A) and may reflect a favouring of the inward-facing state resulting from perturbed binding of cholesterol in the non-flipped orientation (21). The remaining residues characterised would be expected to coordinate the isooctyl tail of cholesterol and, due to flexibility in this region, were not anticipated to alter transport functionality. This was true of the TM5 mutants Y430F/L, however, the TM7 mutants T512A and F515V/W significantly altered glycine transport. T512A generated a 4.8-fold reduction in apparent glycine affinity exhibiting a glycine EC_50_ of 90 μM (p < 0.0001). Due to its distance from cholesterol in the non-flipped orientations it is unlikely that this effect results from disrupting a direct interaction. An indirect effect on cholesterol binding in CHOL1 has previously been proposed for the corresponding residue in hSERT (T371) (*17*). In hSERT, T371 interacts with both V367 (T508 in GlyT2) and Y289 (Y430 in GlyT2), a residue our simulations suggest is critical in mediating cholesterol binding in GlyT2. Both the T371A and V367N mutants resulted in more inward-facing conformations of hSERT suggesting removal of this interaction perturbs cholesterol binding (*17*). Furthermore, Laursen *et al*. (17) suggest that the hSERT T371A mutation destabilises the conformation of Y289 (T508 in GlyT2) (*17*). Thus, the GlyT2 T512A mutation presented here may disrupt intramolecular interactions with both T508 and Y430, perturbing binding of non-flipped cholesterol through a similar mechanism and altering the transporter conformation such that there is a shift in apparent glycine affinity. In contrast to T512A, F515V and F515W enhanced apparent glycine affinity, shifting the EC_50_ to 7.8 μM (p = 0.0147) and 7.6 μM (p = 0.0034), respectively. As F515 is located one helical turn above T512, it may also be disrupting cholesterol binding via an indirect mechanism, altering the glycine EC_50_.

**Figure 8.**
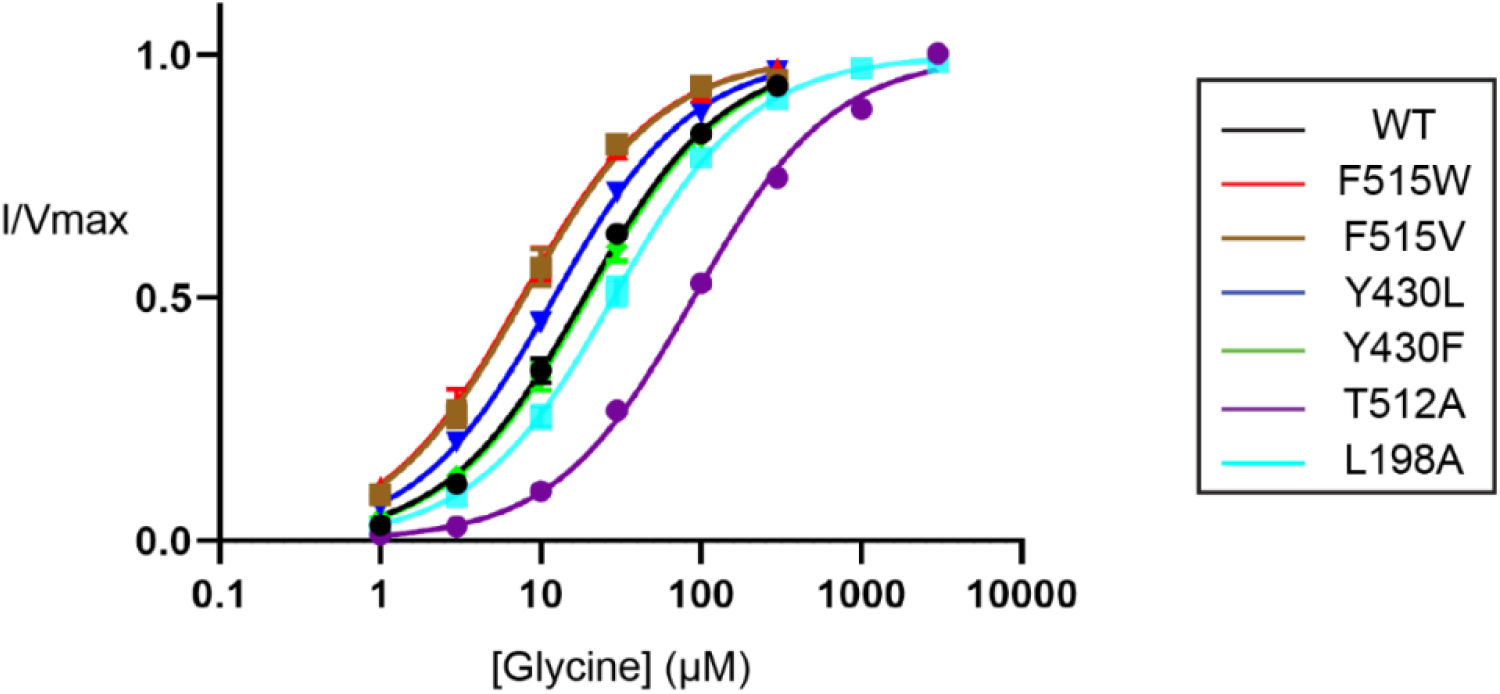
Glycine-dependent currents of WT and CHOL1 mutant GlyT2 transporters expressed in *Xenopus laevis* oocytes. The transport activity of WT and mutant GlyT2 transporters was determined by measuring glycine-dependent currents in response to increasing concentrations of glycine from 1 μM to 3 mM. Currents were fit to the modified Michaelis-Menten equation (see Methods), and raw currents were normalised to the V_max_ for each cell. Data points are presented as mean ± SEM with n ≥ 3 from at least two batches of oocytes.

**Figure 9.**
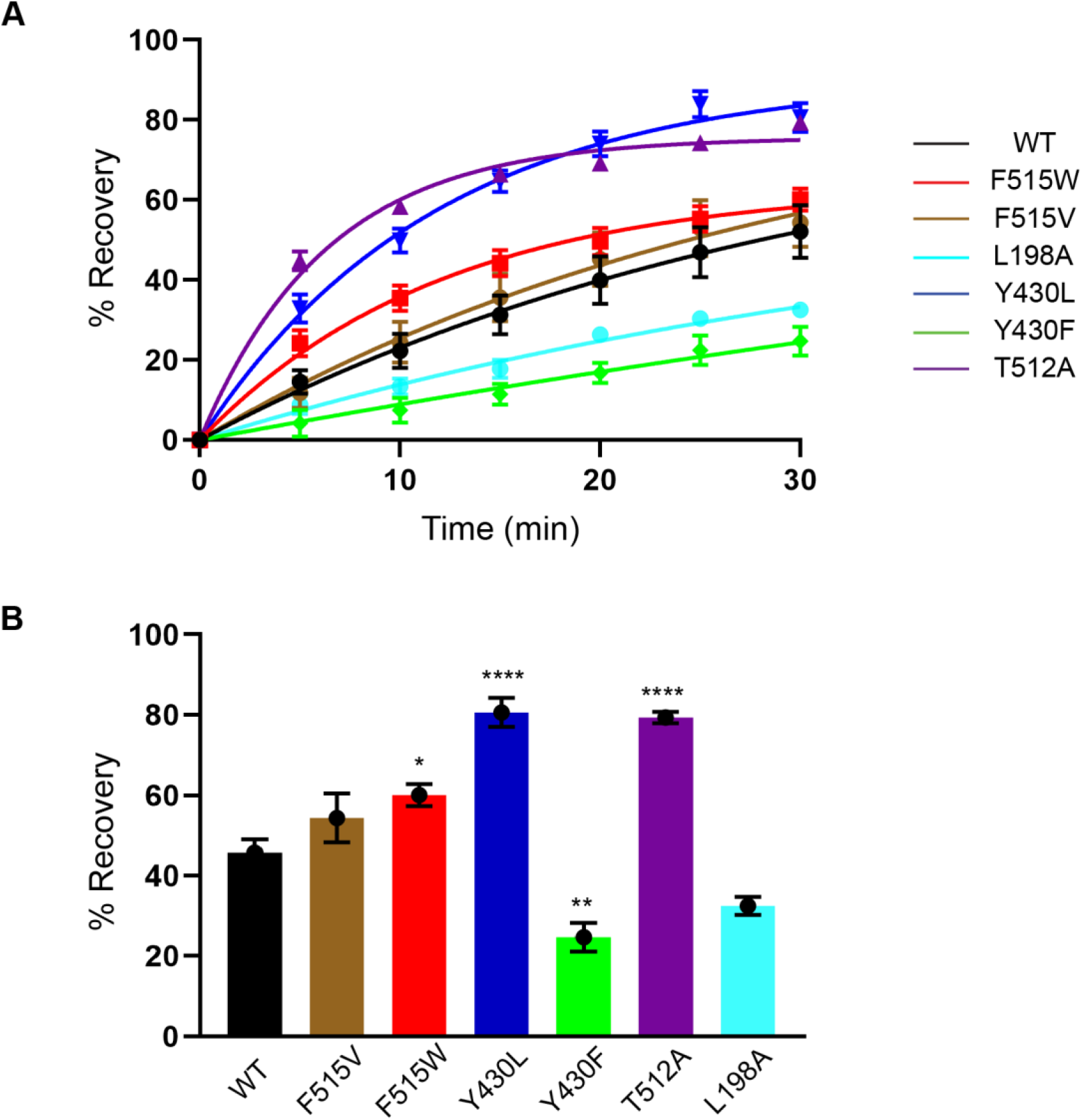
Reversibility of Oleyol-L-Carnitine inhibition of WT and CHOL1 mutant GlyT2 transporters expressed in Xenopus laevis oocytes. OLCarnitine reversibility time course assays (**A**) and recovery of current after 30 minutes (**B**) are shown. Reversibility was determined by co-applying 1 μM OLCarnitine with an EC_50_ concentration of glycine to *Xenopus laevis* oocytes expressing WT and mutant GlyT2 transporters for 4 minutes. Following exposure to OLCarnitine, the EC_50_ of glycine was re-applied at 5-minute intervals for 30 minutes. Raw currents were normalised to currents generated by application of the glycine EC_50_ alone. Data points are presented as mean ± SEM with n ≥ 3 from at least two batches of oocytes. Differences in recovery between WT and mutant transporters were determined via a one-way ANOVA with a Tukey’s posthoc test. Statistical significance is presented as * p ≤ 0.05, ** p ≤ 0.01, *** p ≤ 0.001 and **** p ≤ 0.0001.

The inhibitory activity of bioactive lipids was examined by measuring concentration-responses on CHOL1 mutant transporters. Consistent with observations under cholesterol depletion conditions, the inhibitory activity of OLCarn was unaffected by the majority of CHOL1 mutants except Y430F (Figure 10, Table EV10). Application of 1 μM OLCarn produced significantly greater inhibition of Y430F compared to WT, increasing the level of inhibition by 15.5% (p = 0.0156). Although no differences in IC_50_ values were found for these transporters, the increased level of inhibition likely reflects an increase in potency rather than efficacy as these curves do not plateau, limiting determination of the IC_50_. More refined IC_50_ values could not be determined due to membrane disruptive effects observed with higher concentrations of OLCarn preventing their characterisation. Changes in the inhibitory activity of other bioactive lipids also aligned well with the cholesterol depletion experiments. The activity of OLTrp, which is not predicted to interact with cholesterol and was unaffected by cholesterol depletion, was not altered by any of the mutations (Figure 10, Table EV10). In contrast, OLLys and OLLeu, both of which had reduced potency following cholesterol depletion, were affected by CHOL1 mutants (Figure 10, Table EV10). The potency of OLLys was reduced 4-fold from 220 nM on WT to 860 nM on Y430L (p = 0.0407). Additionally, the inhibition at 3 μM was reduced by 16.9% (p = 0.0148) indicating that the actual shift in potency is likely to be larger if higher concentrations of the compound could be tested. The potency of both OLLys and OLLeu was shifted by the T512A mutation as inferred by reductions in the level of inhibition by 18.6% (p = 0.0073) and 35.9% (p = 0.0053), respectively. In contrast to the T512A mutation, application of 3 μM OLLeu produced significantly greater inhibition of F515W, increasing by 17.3% (p = 0.0098), suggesting an increase in potency.

**Figure 10.**
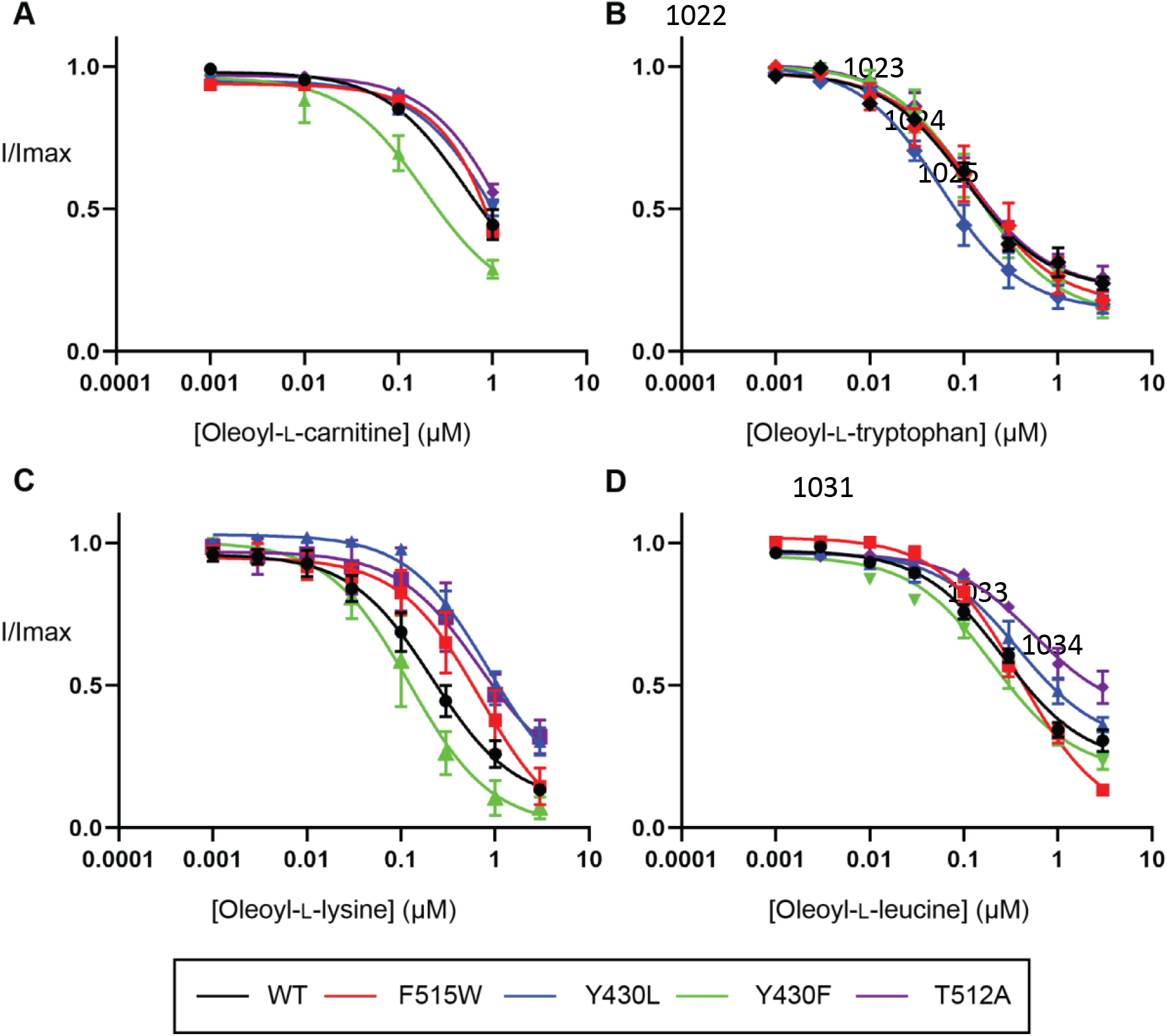
Inhibitory activity of bioactive lipids at WT and CHOL1 mutant GlyT2 transporters expressed in Xenopus laevis oocytes. An EC_50_ concentration of glycine was co-applied with increasing concentrations of (**A**) Oleyol-L-Carnitine, (**B**) Oleyol-L-Tryptophan, (**C**) Oleyol-L-Lysine, and (**D**) Oleyol-L-Leucine ranging from 1 nM to 3 μM. Raw currents were normalised to currents generated by application of the glycine EC_50_ alone. Data points are presented as mean ± SEM with n ≥ 5 from at least two batches of oocytes.

To characterise the effect of CHOL1 mutants on the reversibility of bioactive lipids, washout assays were performed as described above. No recovery of current was observed following application of OLTrp to any of the CHOL1 mutants (Figure 11 and Table EV11), consistent with the observation that membrane cholesterol does not modulate the activity of this bioactive lipid. In contrast, the reversibility of OLCarn, OLLeu and OLLys were significantly altered by the CHOL1 mutations. The reversibility of OLCarn was enhanced with T512A, compared to WT GlyT2, with recovery after 30 minutes increased 1.8-fold from 45.7% to 79.4% (p < 0.0001). F515W also significantly increased the recovery of OLCarn at 30 minutes by 1.3-fold to 60.1% (p = 0.0432) whilst it was unaltered by F515V. Intriguingly, the TM5 mutants Y430F/L had opposing effects on the recovery of OLCarn. Reducing the steric bulk in the region of Y430 via mutation to a leucine (Y430L) resulted in significantly enhanced recovery of OLCarn after 30 minutes to 80.6% (p < 0.0001). In contrast, conserving the steric bulk but removing the polarity of the hydroxyl group via mutation to a phenylalanine (Y430F) produced a significant reduction in reversibility of OLCarn with only 24.7% (p = 0.0002) recovery of current achieved after 30 minutes. This result highlights the potential significance of the aromatic ring mediating interactions with bound cholesterol. Changes in OLLeu reversibility aligned well with what was observed for OLCarn (Figure 11 and Table EV11). Specifically, F515W, Y430L and T512A increased the recovery of current 30 minutes after OLLeu application by 19.0% (p = 0.0001), 16.8% (p = 0.0004) and 25.1% (p < 0.0001), respectively; while Y430F reduced recovery by 24.5% (p < 0.0001). In addition, as the recovery from OLLeu inhibition reached a stable plateau on each of the transporters tested, the half-life was also able to be determined. No changes were observed in the half-life recovery of OLLeu on any of the mutants (Table 3) suggesting a disrupted cholesterol interaction increases the total recovery but does not alter the rate at which that recovery is achieved. Lastly, both the Y430L and Y430F mutations increased the recovery of OLLys after 30 minutes by 17.4% (p = 0.0004) and 13.4% (p = 0.0044) respectively (Figure 11 and Table EV11). The enhanced reversibility of OLLys at Y430F contrasts with observations for other lipids and may result from the altered interactions between Y430F and cholesterol due to changes in cholesterol orientation in the presence of OLLys.

**Figure 11.**
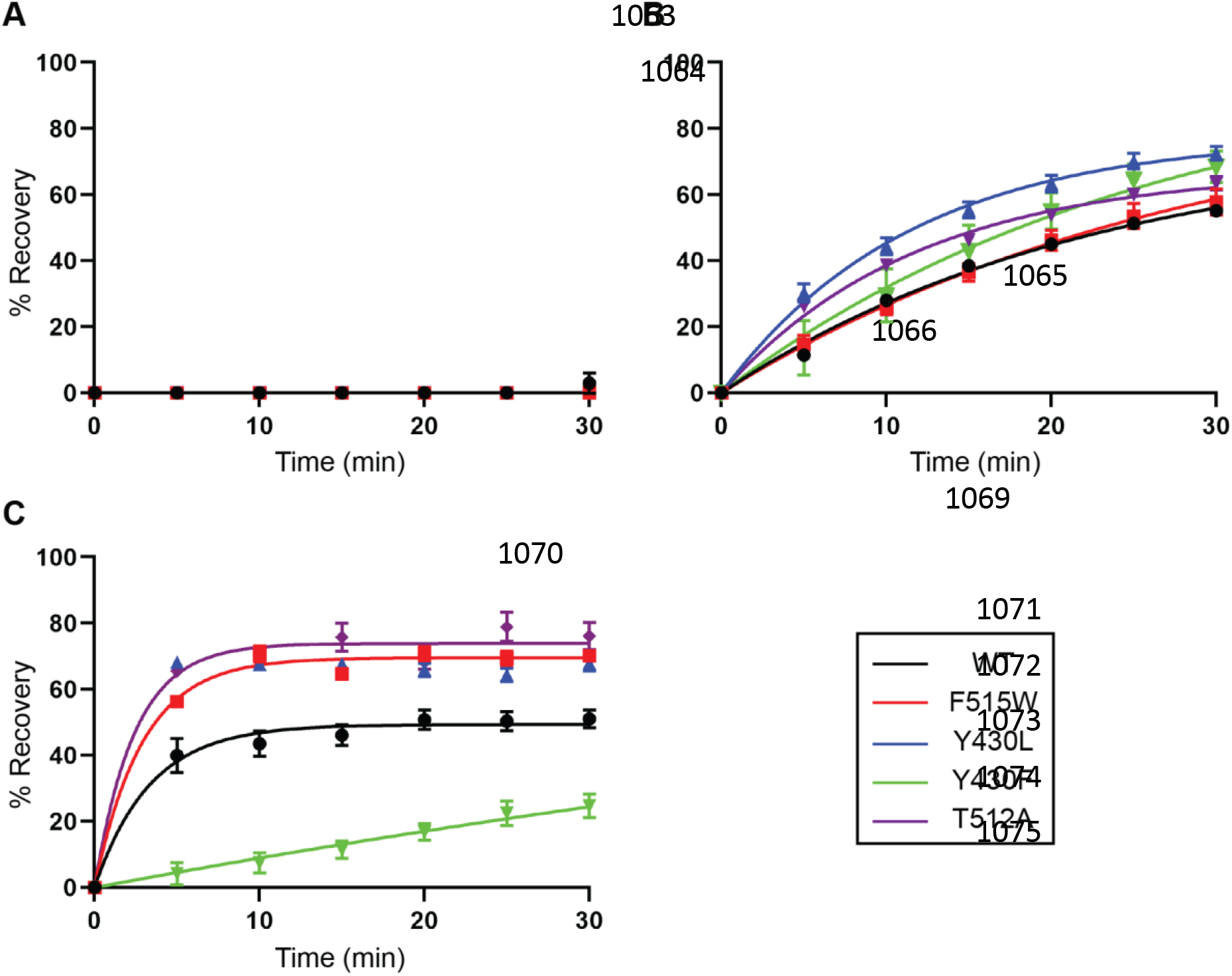
Reversibility of bioactive lipid inhibition of WT and CHOL1 mutant GlyT2 transporters expressed in Xenopus laevis oocytes. Washout time course of (**A**) Oleyol-L-Tryptophan, (**B**) Oleyol-L-Lysine, (**C**) Oleyol-L-Leucine. Washouts were performed by co-applying an IC_50_ concentration of inhibitor with an EC_50_ concentration of glycine to *Xenopus laevis* oocytes expressing WT and mutant GlyT2 transporters for 4 minutes. Following exposure to the inhibitors, the EC_50_ of glycine was reapplied at 5-minute intervals for 30 minutes. Raw currents were normalised to currents generated by application of the glycine EC_50_ alone. Data points are presented as mean ± SEM (n ≥ 5).

To understand the molecular basis for the effect of GlyT2 mutants on inhibitor activity (F515W, T512A, Y430F and Y430L), triplicate 500 ns atomistic molecular dynamics simulations were performed with OLCarn bound to the LAS and a cholesterol molecule in the CHOL1 binding site. Regardless of the mutation, little change is observed in the positioning of OLCarn in the LAS (Figure 12 and Table EV12), however there were changes in the interaction of bound cholesterol and the nearby residues forming the boundary between the LAS and CHOL1. In the T512A mutant, the loss of steric bulk increased the mobility of bound cholesterol but did not significantly change the distribution of residues that interact with cholesterol (i.e., V205, Y430, F515 A208 and V434; Figure 12 and Figure 12 Supplement Table 2). In the combined 1500 ns simulation of F515W, cholesterol remains bound for a greater portion of the total simulation time than observed for the WT system (86% vs 66%, respectively). In one replicate of the F515W system, the bound cholesterol moved out of the binding site, such that the polar head of cholesterol interacted with the heterocyclic amine of F515W. When cholesterol exited the CHOL1 binding site, a simultaneous rotation of M570 blocked the hydrophobic interactions between cholesterol and the lipid inhibitor. Contacts between the CHOL1 residue V205 and Y430 were maintained (Figure 12).

**Figure 12:**
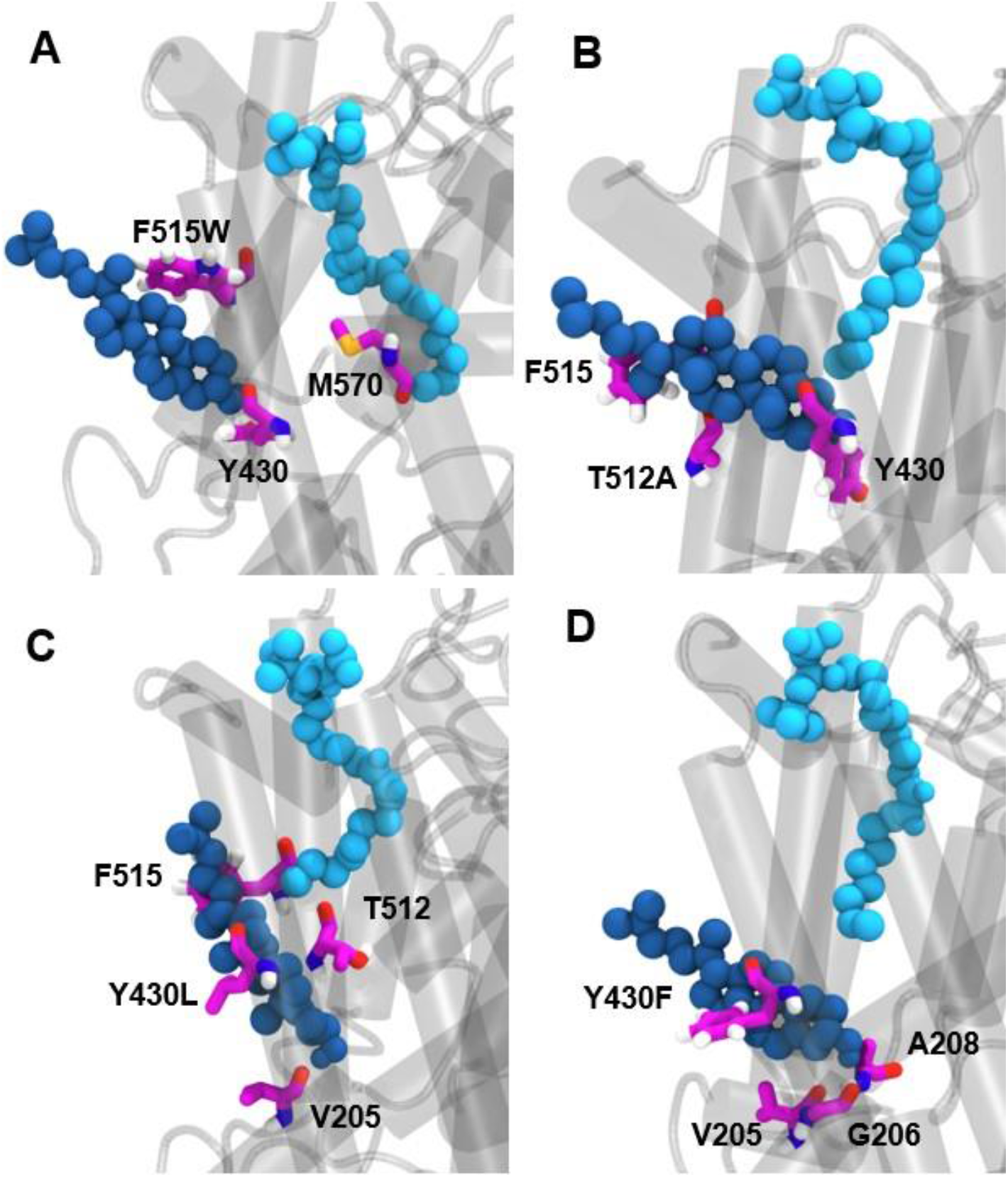
Representative molecular dynamics snapshot of the interaction between bound cholesterol, CHOL1 binding site residues, and Oleyol-L-Carnitine in the four GlyT2 CHOL1 site mutants. Oleyol-L-Carnitine (skyblue) occupies the LAS in simulations of the a) F515W b) T512A c) Y430L d) Y430F GlyT2 mutant. Cholesterol is in dark blue and key CHOL1 residues are fuchsia.

In WT GlyT2 simulations, cholesterol interacted with Y430 for 70% of the total simulation time. When Y430 is mutated to leucine (Y430L), the total contact time is reduced to 63%. The reduced contact time can be attributed to the loss of steric bulk on mutation to leucine which reduced the coordination of cholesterol within the cavity and significantly increased its mobility. In the Y430L simulations, cholesterol reoriented to align parallel to OLCarn (Figure 12). The reoriented cholesterol maintains contact with T512, F515 and V205 observed in the WT simulations, and forms new contacts with G206, Y207, and L433. In contrast, the Y430F mutation increased hydrophobicity while maintaining steric bulk. Cholesterol maintained contact with Y430F for 98% of the trajectory, pointing to the significance of the aromatic interaction. Cholesterol was observed to bind deeper into the binding pocket (Figure 12), forming new contacts with G206 and A208 near the conserved V205 contact. Similarly, contacts were formed with V434 near F430, A511 and T512 near F515. This subsequently increased the hydrophobicity and stability of the OLCarn binding site, reducing the repulsion of the alkyl tail.

## Discussion

In previous studies we have shown that bioactive lipids bind to the LAS of GlyT2, and act as inhibitors of substrate transport (*23*, *27*). Here we show that the allosteric inhibition of substrate transport by lipid inhibitors at the GlyT2 LAS is modulated by the recruitment of membrane cholesterol from the vicinity of the CHOL1 site. The recruitment mechanism involves the flipping of cholesterol from its orientation in the membrane, the subsequent insertion of the cholesterol 3’ hydroxyl group into a cholesterol binding cavity formed between TM1, 5 and 7, followed by its interaction with the base of the LAS and bound inhibitor. The sensitivity and potency of inhibition corresponds to the interaction of the cholesterol 3’ hydroxyl group with the acyl chain terminus of the lipid inhibitor. Of the four different lipid inhibitors investigated, inhibitors that protruded deeper into the LAS, altered the relative orientation of the Y430 (TM5) side chain to create a cavity that membrane cholesterol occupying the CHOL1 site enters and binds to. We show that depletion of membrane cholesterol, and mutation of the residues in the CHOL1 site, also reduces the efficacy of lipid inhibitors bound to the LAS, supporting the cholesterol-modulation of inhibitor activity. Previous studies have shown that application of the cholesterol sequestering agent, mβCD, alters the activity and reversibility of the GlyT2 lipid inhibitor OLCarn (*28*). While this effect was originally attributed to a direct interaction between mβCD and OLCarn, the potential role of cholesterol was not explored. We propose that the binding of cholesterol to the CHOL1 site, stabilises the outward-facing conformation of GlyT2 to facilitate the formation of an accessible LAS and the subsequent binding of lipid inhibitors. Our coarse-grained MD simulations indicate that the lipid inhibitors access the LAS directly from the extracellular solution, rather than from the membrane. Additionally, once in the LAS, the mechanism of inhibition appears to be modulated by cholesterol from the CHOL1 site binding between TM1, TM5 and TM7. We hypothesize that the deep penetration of lipid inhibitors into the LAS is critical for the interaction of cholesterol with the lipid inhibitor, which together may further stabilise GlyT2 in an outward open conformational state, inhibiting the transport process. Therefore, the more pronounced effect of cholesterol depletion on the cholesterol-dependent inhibitors OLLys and OLCarn, compared to the cholesterol-independent inhibitor OLTrp may be due to the deeper penetration of the lipid tail which facilitates a direct interaction between the lipid inhibitor and cholesterol.

Cholesterol has long been implicated in the regulation of the neurotransmitter/sodium symporters (NSSs) of the SLC6 transporter family, however until recently the mechanism has been largely unknown (*4*). Recent structural data, simulations and experimental studies have identified five conserved cholesterol binding sites in the outward open conformations of homologous NSS transporters (*7*, *17*, *18*). Of these cholesterol binding sites, binding of cholesterol to the CHOL1 site has been shown to shift the transporter conformational equilibrium, stabilising the outward open conformation of DAT and SERT (*7*, *17*).Furthermore, membrane cholesterol depletion has been shown to decrease the activity of modulators that bind the outward open conformation and increase the activity of modulators that bind to the inward open conformation (*4*, *17*), supporting the proposed role of cholesterol in stabilising the outward open conformation.

Cholesterol is heterogeneously distributed in neuronal cells and constitutes 44.6 mol % of the neuronal membrane (*18*, *29*). Plasma cholesterol is transported in lipoproteins, which cannot cross the blood brain barrier. As a result, cholesterol homeostasis is independent of plasma cholesterol levels and neuronal cholesterol is synthesized *in situ* (*30*) and its concentrations are closely regulated in normal brain function (*30*). Defects in brain cholesterol metabolism and homeostasis has been shown to be implicated in psychiatric and neurodegenerative diseases (*30*). Because neuronal cholesterol is synthesized *in situ*,cholesterol lowering agents such as statins impact neuronal cholesterol biosynthesis (*31*). Further studies have shown that statins can be detected at significant levels in the brain following a single dose (*31*, *32*), and that long-term treatment with the statin simvastatin reduces the cholesterol content of membrane lipid rafts and significantly reduces cholesterol levels in the extracellular leaflet of brain cells, altering membrane properties such as fluidity (*33*). These observations, together with the results in this study, suggests that the actions of brain penetrating statins may influence glycine transport and reduce the efficacy of glycine transport inhibitors. Thus, the future development of GlyT2 inhibitors for the treatment of neuropathic pain may need to consider their effectiveness in patients being treated with statins.

Modulation of neurotransmitter transporter function underpins the management of a number of neurological conditions, including depression, anxiety, epilepsy, Parkinson’s disease and addiction, as well as chronic pain (*34*). When considered together, the previous data for SERT, DAT (*7*, *17*), as well as the results for GlyT2 presented here, strongly suggest that variations in membrane cholesterol levels influence the conformational equilibria of neurotransmitter transporters during the transport cycle, and therefore, the physiological clearance of neurotransmitters from the synaptic cleft. Furthermore, our results indicate that membrane cholesterol may impact the activity and efficacy of inhibitors of GlyT2 and other homologous neurotransmitter transporters. Specifically, depletion of cholesterol from membrane lipid rafts may reduce the binding of cholesterol to the CHOL1 site, shifting the conformational equilibrium to the inward open conformation reducing the accessibility of both the substrate binding site and LAS. A failure to respond to SSRIs, SNRIs and competitive inhibitors of neurotransmitter transporters is notable in some patients, one of the factors that may contribute to the lack of efficacy could be related to membrane cholesterol regulation.

The interaction of cholesterol in the CHOL1 site and bound inhibitors in the LAS provides a promising and novel avenue for optimisation of pharmaceutical designs that will promote both a selective and reversible inhibition of GlyT2 to treat chronic pain. The observations that cholesterol can influence the equilibrium between inward and outward facing states, suggests that the process could also be pharmacologically manipulated by derivatives of cholesterol – such as the various neurosteroids (allopregnenalone), which would be analogous to the neurosteroid manipulation of ligand-gated ion channels. Drugs may be developed that mimic the effects of cholesterol. Furthermore, chronic pain is associated with increased levels of reactive oxygen species and oxidative stress (*35*, *36*). Cholesterol is more susceptible to oxidation by reactive oxygen species than phospholipids due to the 6-double bond and vinylic methylene group at C-7 in the B ring of the sterene, as well as the isooctyl side chain at C-17 (*37*), and oxysterols are important regulators of neuronal lipid homeostasis (*30*). As neuronal membrane oxysterol levels can increase to up to 20 mol % in pathological conditions, competitive binding between oxysterols and cholesterol at the GlyT2 CHOL1 site must also be considered in a personalised medicine approach for pharmacological design of GlyT2 inhibitors of the ascending pain pathway.

Another implication of the results presented in this study is whether a similar allosteric ligand binding site exist in other closely related SLC6 transporters for dopamine, serotonin, and GABA. If this site is present on these other transporters, it may provide the opportunity to develop new classes of neurotransmitter transport inhibitors for the treatment of a range of neurological disorders such as depression, drug addiction, Parkinsons Disease and schizophrenia.

## Materials and Methods

### Molecular Dynamics

#### Coarse Grain Simulations

The GROMACS 2016.1 molecular dynamics engine^51^ and the MARTINI2.2P forcefield (*38*) were used for all coarse-grained simulations. The previously published homology model of GlyT2 with 2 Na^+^ ions and the substrate glycine bound was used as a starting structure for modeling (*39*, *40*). GlyT2 was converted to a GoMARTINI coarse-grained model using the *go_martinize* package with Gō contacts of 9.414 kJ/mol.^40^ The substrate ions were manually converted to coarse-grain MARTINI 2.0 ions. The protein and its substrates were embedded in an 80% POPC and 20% cholesterol (CHOL) bilayer and solvated with polarizable (martini 2.2P) coarse grain water (box size: 166 × 194 × 160 Å) using insane.py (*41*). The bilayer was modeled with MARTINI 2.0 lipids. The system was solvated with the MARTINI polarizable water model.^54^ To replicate the patch-clamping experiments where the additives are only applied to the extracellular side of the cell and simulate spontaneous binding of the lipid inhibitors to the extracellular leaflet, a layer of position restrained water was created by selecting a single later of water molecules approximately 10 Å below the intracellular leaflet. The layer of position restrained water was extended in the x and y directions to ~1.5 Å from the box edges to allow for changes in the box size upon simulation. The water in this layer was restrained with a force constant of 1000 kJ mol^-1^ nm^-2^ for the duration of the simulations. Twenty lipid inhibitors of a single type, namely OLLys, OLLeu, OLSer or OLTrp, were placed at random coordinates within the extracellular solution or between the layer of the position restrained water and box edge. The systems were neutralized and NaCl was added to a concentration of 150 mM to mimic physiological conditions.

The entire system was minimized and then equilibrated with a force constraint on the protein backbone. The constraint on the protein backbone was slowly reduced through five sequential simulations of 1 ns each, with constraint weights of 1000, 500, 100, 50 and 10 kJ mol^-1^ nm^-2^, respectively. Each system was then simulated for 10 μs in triplicate without constraints on the protein. A new random starting velocity was assigned at the start of each replicate simulation. Coordinates were saved every 20 ns, and a time step of 5 and 20 fs were used for the equilibration and production simulations, respectively. All simulations were performed in the NPT ensemble. The simulations were performed at 310 K and 1 bar with the velocity rescale thermostat (τ_T_=1 K) and Berendsen barostat (τ_p_=3.0 bar). The pressure was modeled with semi-isotropic pressure coupling (isothermal compressibility=3.0×10^-4^ bar^-1^), with the pressure being isotropic in the plane of the bilayer. The length of covalent bonds were constrained using the LINCS algorithm. Electrostatic interactions were calculated using the reaction-field method and van der Waals interactions were calculated with a cut-off of 11 Å.

#### Backmapped Atomistic Simulations

Atomistic simulations were performed using the GROMACS 2016.1 MD package (*42*) and the GROMOS 54a7 forcefield (*26*). Lipid inhibitors were modeled using parameters generated using the Automated Topology Builder and Repository (ATB) (*43*), (OLLys MoleculeID: 252919, OLTrp MoleculeID: 252930, OLLeu MoleculeID: 252921, and OLCarn MoleculeID: 296337) as previously described (*27*). To ensure that there was no isomerisation around the cis double bond, the force constant related to this dihedral angle was adjusted from 5.86 kJ/mol/rad^2^ to 41.80 kJ/mol/rad^2^, as previously described (*27*). The water model was SPC (*44*). To further understand the binding of the lipid inhibitors to GlyT2, selected frames from simulations in which OLLeu was bound at the LAS and CHOL was interacting with the CHOL1 site were backmapped to atomistic coordinates using backmap.py (*45*). Only lipid inhibitors within 10 Å of the protein were retained in atomistic simulations. All water was treated as unrestrained. This local region was then embedded within an 80% POPC/20% CHOL bilayer and solvated such that the final system size was 166 x 194 x 100 Å. The system was minimized and then equilibrated with a harmonic constraint on the protein CA atoms. The harmonic constraints on the CA atoms were slowly reduced over five sequential 1 ns simulations from 1000, to 500, 100, 50 and finally 10 kJ mol^-1^ nm^-2^. Each system was then simulated for 500 ns in triplicate without constraints with a 2 fs timestep. A new random starting velocity was assigned at the start of each replicate simulation. Coordinates were saved every 0.1 ns. All simulations were performed in the NPT ensemble. The simulations were performed at 310 K and 1 bar with the velocity rescale thermostat (τ_T_=0.1 K) and Berendsen barostat (τ_p_=0.5 bar). The pressure was modeled with semi-isotropic pressure coupling (isothermal compressibility=4.5×10^-5^ bar^-1^), with the pressure being isotropic in the plane of the bilayer. The length of the covalent bonds were constrained using the LINCS algorithm. Electrostatic interactions were calculated using the particle mesh Ewald and van der Waals interactions were calculated with a cut-off of 10 Å.

#### Atomistic Simulations of GlyT2 with docked inhibitor

The equilibrated backmapped system was used as a starting point for the atomistic simulations with OLLys, OLCarn and OLTrp bound to the LAS. OLLeu, solvent and ions were removed from the backmapped simulation. The conformation of the lipid inhibitor in the LAS was obtained by overlaying the final conformation obtained from previous simulations performed with the lipid inhibitor docked into the LAS (*23*). Each system was then solvated, energy minimised, equilibrated and simulated using the protocol outlined above for the backmapped atomistic simulations.

#### Atomistic Simulations of GlyT2 mutations

The equilibrated backmapped system was again used as a starting point for atomistic simulations incorporating the effect of GlyT2 mutations on inhibitor and CHOL binding. Pymol (version 2.5.0, The PyMOL Molecular Graphics System, Schrödinger, LLC) was used to incorporate a series of individual GlyT2 point mutations (T512A, Y430L, Y430F and F515W). To characterise the effect of each mutation on inhibitor and CHOL binding, only one mutation was included per simulation system. Each system was then solvated, energy minimised, equilibrated and simulated using the protocol outlined above for the backmapped atomistic simulations.

#### Atomistic Simulations of inward facing GlyT2

A homology model of the inward facing conformation of GlyT2 was generated using SWISS-MODEL (*46*). Here the inward-facing cryoEM structure of human SERT (PDB ID: 6DZZ) (*25*) was used as a template. The protein was embedded in an 80% POPC and 20% cholesterol bilayer. The inward facing homology model was then solvated, energy minimised, equilibrated and simulated using the protocol outlined above for the backmapped atomistic simulations.

#### Simulation Analysis

Area per lipid and bilayer thickness were calculated using FATSLiM (*47*). Simulations were visualised and contact residues calculated using VMD 1.9.3 (*48*). The pocket volume for the LAS was calculated using trj_cavity (*49*).

Simulation parameter files, and the initial and final coordinates of the simulations are available at https://github.com/OMaraLab/GlyT2_mutants

### Cholesterol depletion and electrophysiology

#### Generation of wild-type (WT) and mutant RNA encoding GlyTs

GlyT2a (referred to herein as GlyT2) was subcloned into the plasmid oocyte transcription vector (pOTV) for expression in *Xenopus* laevis oocytes. Oligonucleotide primers incorporating desired mutations were synthesised by Sigma Aldrich (Sydney, Australia) and used to generate point mutations in the GlyT2 cDNA/pOTV construct utilising the Q5 site-directed mutagenesis kit (New England Biolabs (Genesearch), Arundel, Australia). Amplified PCR products were transformed into chemically competent *E. coli* cells and purified using the GeneJET plasmid miniprep kit (Thermo Fisher Scientific, Sydney, Australia). Purified plasmid DNA was sequence verified by the Australian Genome Research Facility (Sydney, Australia). pOTV constructs were linearised with the restriction enzyme *SpeI* (New England Biolabs (Genesearch), Arundel, Australia) and RNA transcribed by T7 RNA polymerase using the mMESSAGE mMACHINE T7 kit (Thermo Fisher Scientific, Sydney, Australia).

#### Oocyte harvesting and preparation

*Xenopus laevis* were anaesthetised with 0.17% (w/v) 3-aminobenzoic ethyl ester (tricaine). A single incision was made in the abdomen and a lobe of oocytes removed. Individual oocytes were separated from the follicle via digestion with 3 mg/mL collagenase A (Boehringer, Germany) at 18°C for 45-90 minutes. All surgeries were conducted in accordance with the *Australian Code of Practice for the Care and Use of Animals for Scientific Purposes*.

Separated stage IV/V oocytes were injected with 23-4 nL of WT or mutant RNA (Drummond Nanoinject, Drummond Scientific Co., Pennsylvania, USA). Injected oocytes were stored in frog Ringer’s solution (ND96; 96 mM NaCl, 2 mM KCl, 1 mM MgCl_2_. 1.8 mM CaCl_2_, 5 mM HEPES, pH 7.5) supplemented with 2.5 mM sodium pyruvate, 0.5 mM theophylline, 50 μg/mL gentamicin and 100 μM/mL tetracycline. Oocytes were stored at 18°C for 2-4 days until sufficient transporter levels were reached to measure transport activity. Sufficient transporter expression was defined as the onset of robust and readily reproducible inward currents in response to substrate.

#### Two-electrode voltage clamp electrophysiology

The transport activity of GlyT2 is an electrogenic process as it couples the transport of glycine to the co-transport of 3 Na^+^ and 1 Cl^-^ and thus can be measured using two-electrode voltage clamp electrophysiology. Oocytes expressing GlyTs were voltage clamped at −60 mV and glycine induced whole-cell currents were measured using a Geneclamp 500 amplifier (Axon Instruments, California, USA) and a Powerlab 2/26 chart recorder (ADI Instruments, Sydney, Australia) and LabChart Software (ADI Instrument, Sydney, Australia).

#### Membrane cholesterol depletion

Oocyte membranes were depleted of cholesterol by treating them with the cholesterol sequestering agent methyl-β-cyclodextrin (mβCD). Individual oocytes were incubated at 32°C for 30 minutes in 15 mM mβCD, suspended in frog Ringer’s solution. Oocytes were then washed in ND96 for 5 minutes unclamped and a further 5 minutes voltage clamped to ensure complete removal of cyclodextrin.

#### Glycine concentration responses

Functionality of WT and mutant GlyT2 transporters was assessed by applying increasing concentrations of glycine (1-3000 μM) in ND96. EC_50_ values for each transporter were determined by fitting concentration response data to the modified Michaelis-Menten equation:

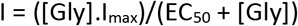

where I is current in nA, [Gly] is the concentration of glycine, I_max_ is the current generated by a maximal concentration of glycine and EC_50_ is the concentration of glycine that generates a half maximal current.

Differences in EC_50_ and V_max_ values between WT and mutant transporters were analysed via a one-way ANOVA with a Tukey’s posthoc test. Statistical significance is presented as * p ≤ 0.05, ** p ≤ 0.01, *** p ≤ 0.001 and **** p ≤ 0.0001.

#### Glycine concentration responses – cholesterol depletion

As glycine is a fully reversible substrate, glycine concentration responses were performed on the same cell pre- and post-treatment with mβCD. Glycine dependent currents following cyclodextrin treatment were normalised to the V_max_ of the control response and then fitted to the modified Michaelis-Menten equation. Differences in EC_50_ and V_max_ values between non-treated and cyclodextrin treated cells were analysed via a paired two-tailed t-test. Statistical significance is presented as * p ≤ 0.05, ** p ≤ 0.01, *** p ≤ 0.001 and **** p ≤ 0.0001.

#### Inhibitor concentration response – mutagenesis

Due to the slowly reversible nature of the compounds tested, inhibitor concentration responses were performed cumulatively. An EC_50_ concentration of glycine was applied until a stable transport current was reached. EC_50_ concentrations of glycine were used to produce reliable currents for each transporter and as the inhibitors are non-competitive, application of varying glycine concentrations is unlikely to affect their activity. Increasing concentrations of inhibitors were co-applied with glycine with each concentration producing a distinct plateau before the subsequent concentration was applied. Currents following the plateau of each inhibitor concentration were normalised to the current produced by application of glycine alone and fit using the method of least squares:

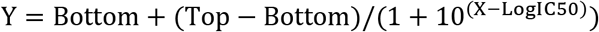

where X is log[inhibitor], Y is current normalised to glycine in absence of inhibitor and Top and Bottom are the maximal and minimal plateaus, respectively. The method of least squares was constrained to have the bottom value > 0 and the hill slope as −1, enabling the calculation of IC_50_ values and the extent of inhibition produced by the maximal concentration applied.

IC_50_ values are presented as mean and 95% confidence interval whereas inhibition values are presented as mean ± SEM. Data sets are n ≥ 5 from two batches of oocytes. For instances where substantial inhibition was not achieved, the IC_50_ value is recorded as greater than the highest concentration of inhibitor applied. Statistical analysis of inhibitor concentration responses across WT and mutant transporters was performed using a one-way ANOVA with a Tukey’s posthoc test for both IC_50_ values and maximal inhibition. Statistical significance is presented as * p ≤ 0.05, ** p ≤ 0.01, *** p ≤ 0.001 and **** p ≤ 0.0001.

#### Inhibitor concentration responses – cholesterol depletion

As the inhibitors assayed here are slowly reversible, inhibitor concentration responses were unable to be performed on the same cell following treatment with MβCD. Thus, GlyT2 expression was tested by applying a maximal glycine concentration (300 μM) before depleting membrane cholesterol. Following washing, 300 μM glycine was re-applied and reduction in V_max_ was used as an indicator of successful depletion. For cholesterol depletion inhibitor concentration responses, an unpaired two-tailed t-test was used to determine differences in IC50 values and the inhibition produced by the maximal concentrations tested, with statistical significance represented as * p ≤ 0.05, ** p ≤ 0.01, *** p ≤ 0.001 and **** p ≤ 0.0001.

#### Inhibitor washout time course

Reversibility of GlyT2 inhibitors was determined by applying an EC_50_ concentration of glycine followed by co-application of the inhibitor at an IC_50_ concentration once a stable current had been achieved. Inhibitors were applied for 4 minutes as this was sufficient time to produce substantial inhibition for each compound and allowed for their direct comparison. At the conclusion of 4 minutes (time 0), ND96 buffer was perfused through the recording chamber to wash the oocyte. At 5-minute intervals, up to 30 minutes, the EC_50_ glycine concentration was re-applied, and recovery of transport used as an indicator of reversibility. Transport currents were normalised to glycine currents in the absence of inhibitor and converted to % recovery values via:

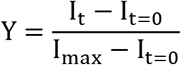

where Y is % recovery, I is current in nA, I_max_ is the maximal current produced by the glycine EC_50_ and t is time in minutes. Washout kinetics were then determined by fitting % recovery values to the first order rate equation:

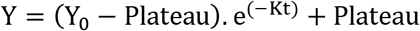

where Y is the % recovery, Y_0_ is the point which crosses the Y axis (constrained to 0) and increases to the Plateau (constrained to between 0 and 100), K is the rate constant (1/t) and t is time in minutes.

Differences in half-life and maximal recovery between WT and mutant transporters were analysed via a one-way ANOVA with a Tukey’s posthoc test. Due to the slowly irreversible nature of the inhibitors, washout time courses involving cyclodextrin treatment could not be performed on the same cell. Thus, differences in half-life and maximal recovery in these experiments were analysed via an unpaired two-tailed t-test. In both instances statistical significance is presented as * p ≤ 0.05, ** p ≤ 0.01, *** p ≤ 0.001 and **** p ≤ 0.0001.

## Acknowledgements

This work was supported by the Australian National Health and Medical Research Council Project Grant APP1144429 to RJV and MLO. Simulations were performed with the assistance of resources and services from the National Computational Infrastructure (NCI), which is supported by the Australian Government.

## Competing interests

The authors declare that they have no conflict of interest.

## Figures and Tables

**Figure EV1.**
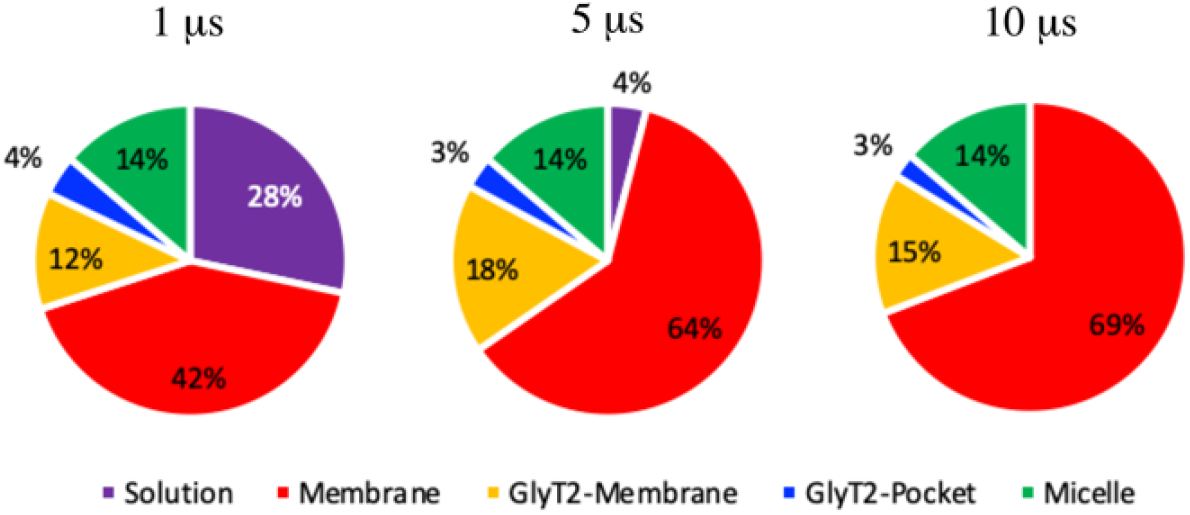
Location of the lipid inhibitors over the course of the CG spontaneous binding simulations, where solution refers to a single lipid inhibitors solvated in the solution environment, micelle refers to a group of lipid inhibitors in the solution environment, membrane refers to the membrane environment and >4 Å from GlyT2, GlyT2-membrane is the lipid inhibitors that are in the membrane and interacting with GlyT2 and GlyT2-pocket is lipid inhibitors that have bound to a solvent accessible pocket of GlyT2.

**Table EV1.**
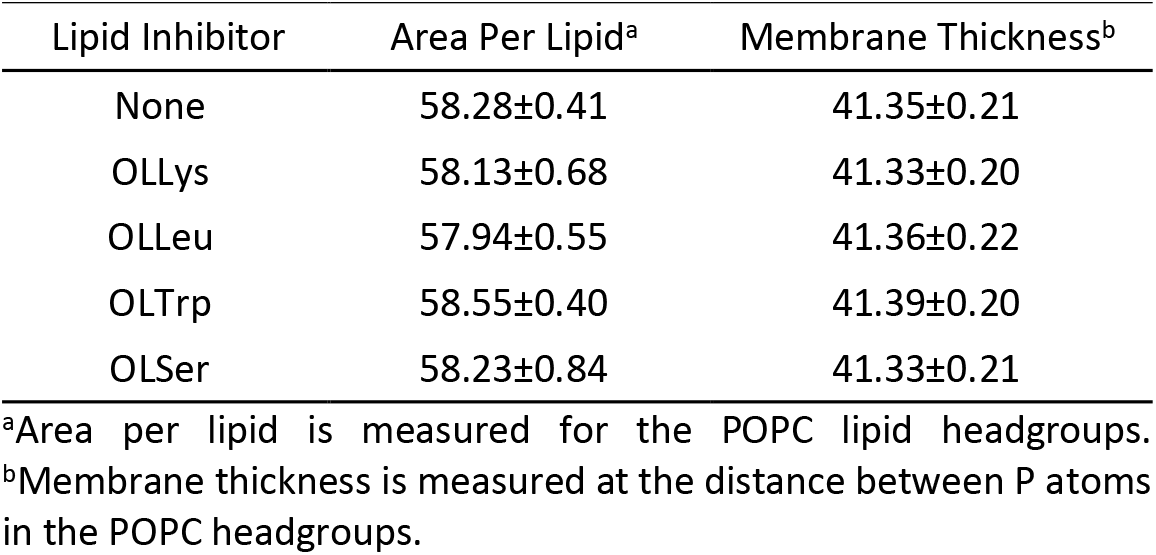
Thickness (Å) and area per lipid (Å^2^) for the lipid bilayer for 10 μs coarse grain simulations on GlyT2 in a POPC/CHOL membrane with no lipid inhibitors (control) or 20 of the specified lipid inhibitors present.

**Table EV2.** Percentage of the total CG simulation time in which residues are in contact with the OLLeu lipid inhibitors that is bound in the extracellular allosteric pocket of GlyT2. Only interactions that occur for >15% of the total simulation time are reported.^a^

| Region | Residue | Occupancy |
| --- | --- | --- |
| TM1 | W215 | 72.5 |
| TM1 | P218 | 48.9 |
| TM1 | Y219 | 46.2 |
| TM5 | L437 | 21.3 |
| TM5 | G440 | 18.1 |
| TM5 | V441 | 34.1 |
| TM7 | V751 | 51.7 |
| TM7 | I520 | 52.8 |
| TM7 | I524 | 86.8 |
| TM7 | F526 | 16.8 |
| TM7 | M527 | 89.5 |
| TM7 | A528 | 44.4 |
| EL4 | R531 | 15.2 |
| EL4 | V533 | 34.2 |
| EL4 | I535 | 42.9 |
| EL4 | V538 | 56.5 |
| EL4 | P543 | 57.7 |
| EL4 | I545 | 34.5 |
| EL4 | A546 | 16.5 |
| EL4 | F547 | 58.3 |
| EL4 | V548 | 46.3 |
| EL4 | Y550 | 60.6 |
| EL4 | A553 | 39.0 |
| EL4 | L554 | 35.8 |
| EL4 | T555 | 17.0 |
| EL4 | R556 | 42.7 |
| EL4 | L557 | 44.0 |
| EL4 | P558 | 36.2 |
| EL4 | L559 | 46.9 |
| EL4 | S560 | 35.4 |
| TM8 | F562 | 26.7 |
| TM8 | W563 | 81.6 |
| TM8 | A564 | 22.2 |
| TM8 | F567 | 83.1 |
| TM8 | F568 | 30.0 |
| TM8 | L571 | 46.9 |
| TM10 | I626 | 15.5 |
| TM10 | Q630 | 58.6 |

<sup>a</sup>An interaction is defined as a minimum distance between beads in the residues to be < 6 Å.

**Table EV3.**
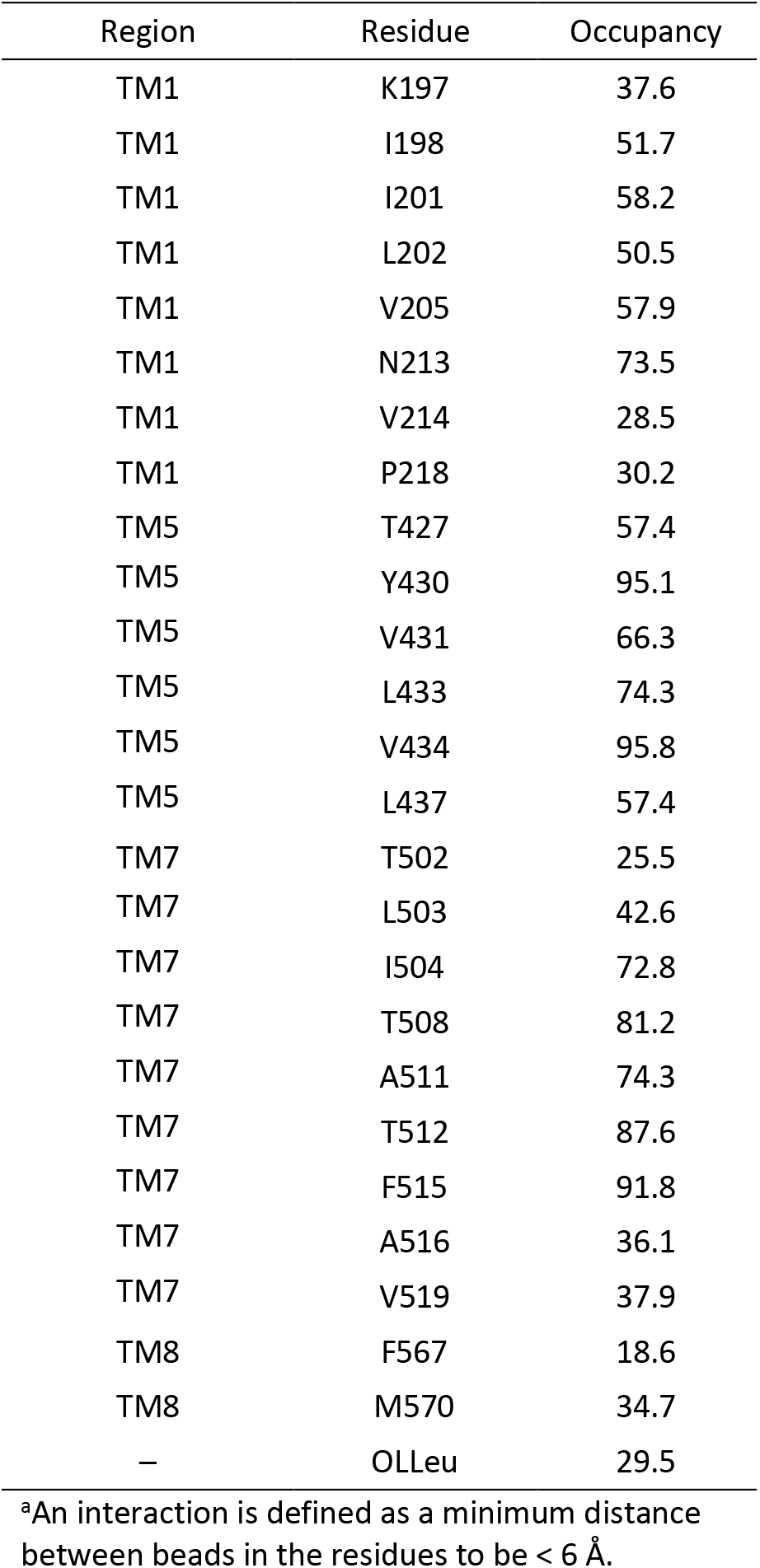
Percentage of the CG simulation time with CHOL interacting in the vicinity of the LAS in which residues are in contact with the cholesterol residues that bound near the bottom of the extracellular allosteric pocket of GlyT2 when OLLeu is bound in the extracellular allosteric pocket. Only interactions that occur for >15% of the total simulation time are reported.^3^

| Region | Residue | Occupancy |
| --- | --- | --- |
| TM1 | K197 | 37.6 |
| TM1 | I198 | 51.7 |
| TM1 | I201 | 58.2 |
| TM1 | L202 | 50.5 |
| TM1 | V205 | 57.9 |
| TM1 | N213 | 73.5 |
| TM1 | V214 | 28.5 |
| TM1 | P218 | 30.2 |
| TM5 | T427 | 57.4 |
| TM5 | Y430 | 95.1 |
| TM5 | V431 | 66.3 |
| TM5 | L433 | 74.3 |
| TM5 | V434 | 95.8 |
| TM5 | L437 | 57.4 |
| TM7 | T502 | 25.5 |
| TM7 | L503 | 42.6 |
| TM7 | I504 | 72.8 |
| TM7 | T508 | 81.2 |
| TM7 | A511 | 74.3 |
| TM7 | T512 | 87.6 |
| TM7 | F515 | 91.8 |
| TM7 | A516 | 36.1 |
| TM7 | V519 | 37.9 |
| TM8 | F567 | 18.6 |
| TM8 | M570 | 34.7 |
| – | OLLeu | 29.5 |

<sup>a</sup>An interaction is defined as a minimum distance between beads in the residues to be < 6 Å.

**Table EV4.** Percentage of the total simulation time in which residues are in contact with the lipid inhibitor bound in the LAS in atomistic simulations of the backmapped structure. Only interactions that occur for >15% of the total simulation time are reported.^a^

| Region | Residue | OLLeu | OLLys | OL-CARN | OLTrp |
| --- | --- | --- | --- | --- | --- |
| TM1 | N213 | 16.3 | 33.3 | - | - |
| TM1 | V214 | 23.9 | 54.7 | 48.6 | 43.9 |
| TM1 | W215 | 29.8 | - | - | - |
| TM1 | F217 | 19.6 | 67.6 | 54.8 | 40.6 |
| TM1 | P218 | 64.6 | 50.7 | 62.3 | 68.0 |
| TM5 | Y430 | 25.8 | 50.5 | 44.9 | 42.7 |
| TM5 | L433 | 26.1 | 42.8 | 24.2 | 40.1 |
| TM5 | V434 | - | 25.3 | 17.8 | 17.0 |
| TM5 | L437 | 50.9 | 77.8 | 53.7 | 74.8 |
| TM5 | G440 | 32.8 | 44.1 | 51.6 | 63.0 |
| TM5 | V441 | 33.2 | - | - | 15.7 |
| TM5 | T442 | 20.4 | - | - | 26.4 |
| TM7 | T512 | 20.1 | 41.9 | 29.3 | 33.0 |
| TM7 | F515 | 22.4 | 43.5 | 58.3 | 66.8 |
| TM7 | A516 | 34.8 | 62.5 | 51.1 | 50.9 |
| TM7 | V519 | 21.2 | 43.1 | 28.2 | 31.3 |
| TM7 | I520 | 57.9 | 66.5 | 60.3 | 71.5 |
| TM7 | V523 | 66.6 | 67.6 | 45.1 | 70.1 |
| TM7 | I524 | 17.4 | - | - | - |
| TM7 | F526 | 36.2 | 37.2 | 39.4 | 56.1 |
| TM7 | M527 | 69.3 | 61.1 | 53.7 | 78.8 |
| EL4 | N529 | - | 20.3 | - | - |
| EL4 | V533 | 19.7 | 48.7 | 75.4 | 47.5 |
| EL4 | I535 | 37.6 | 41.7 | 52.4 | 63.1 |
| EL4 | N537 | 32.0 | 51.5 | 35.9 | 67.5 |
| EL4 | V538 | 42.7 | 19.3 | 38.2 | 60.7 |
| EL4 | G542 | 25.3 | - | - | - |
| EL4 | P543 | 32.3 | - | - | - |
| EL4 | R556 | 26.9 | - | - | - |
| EL4 | P558 | 15.9 | - | - | - |
| EL4 | L559 | 27.3 | 18.5 | - | 61.4 |
| EL4 | S560 | 42.1 | 46.0 | 64.1 | 73.8 |
| TM8 | F562 | 69.2 | 98.3 | 97.4 | 90.5 |
| TM8 | W563 | 100.0 | 94.9 | 88.3 | 97.5 |
| TM8 | I566 | 52.7 | 68.6 | 68.4 | 65.7 |
| TM8 | F567 | 72.8 | 17.7 | 75.6 | 69.8 |
| TM8 | M570 | 66.5 | 73.6 | 74.9 | 69.4 |
| TM8 | L574 | - | 19.3 | - | - |
| TM10 | F629 | - | - | 21.4 | - |
| TM10 | N630 | 22.7 | - | - | - |

<sup>a</sup>An interaction is defined as a minimum distance between heavy atoms in the residues to be < 4 Å.

**Table EV5.** Percentage of the total simulation time (in atomisitic simulations) in which residues are in contact with the cholesterol molecule bound near the bottom of the LAS of GlyT2 while a lipid inhibitor is bound in the extracellular allosteric pocket. Only interactions that occur for >30% of the total simulation time are reported.^a^

| Region | Residue | OLLeu | OLLys | OLCarn | OLTrp |
| --- | --- | --- | --- | --- | --- |
| TM1 | V205 | 37.7 | 52.9 | 60.7 | 55.1 |
| TM1 | G206 | - | - | 35.9 | 35.4 |
| TM1 | A208 | - | 42.5 | 47.3 | 37.5 |
| TM5 | Y430 | 65.3 | 54.5 | 70.4 | 56.4 |
| TM5 | V434 | - | 38.3 | 44.7 | 39.2 |
| TM7 | C507 | 34.3 | - | - | - |
| TM7 | T508 | 32.3 | 32.6 | - | - |
| TM7 | A511 | 43.1 | 31.1 | 43.8 | 38.3 |
| TM7 | T512 | 44.1 | 48.6 | 47.5 | 37.4 |
| TM7 | F515 | 56.3 | 55.7 | 66.1 | 72.6 |
| - | Membrane CHOL | 52.7 | - | - | - |

<sup>a</sup>An interaction is defined as a minimum distance between heavy atoms in the residues to be < 4 Å.

**Table EV6.** Membrane cholesterol depletion alters GlyT2 functionality^†^.

| Treatment | Kinetic Parameters |  |
| --- | --- | --- |
| MβCD | $K_m$ (μM) | $V_{\max}$ |
| Baseline | 18.59<br>(16.44 – 21.02) | 0.9984<br>(0.9666 – 1.031) |
| 0 mM | 15.26<br>(11.23 – 20.63) | 1.041<br>(0.9639 – 1.124) |
| 15 mM | 11.66<br>(8.57 – 15.78) | 0.5936****<br>(0.5508 – 0.6390) |

<sup>†</sup> GlyT2 function was assessed by measuring transport dependent currents following application of increasing glycine concentrations (1-3000 μM) before, and after, incubation of *Xenopus laevis* oocytes with methyl-β-cyclodextrin (mβCD) for 30 minutes at 32°C. Values are presented as mean (95% confidence interval) with $n \geq 5$ . Differences in $K_m$ and $V_{\max}$ values between control and MβCD or $\gamma$ CD treated oocytes were determined using a two-way paired t-test. Statistical significance is presented as \* $p \leq 0.05$ , \*\* $p \leq 0.01$ , \*\*\* $p \leq 0.001$ and \*\*\*\* $p \leq 0.0001$ .

**Table EV7.** Activity of bioactive lipids on WT GlyT2 expressed in control and MβCD treated *Xenopus laevis* oocytes^†^.

| Compound | Condition |  |  |  |
| --- | --- | --- | --- | --- |
|  | Control |  | MβCD |  |
|  | IC<sub>50</sub> (nM) | Inhibition at 3 μM (%) | IC<sub>50</sub> (nM) | Inhibition at 3 μM (%) |
| **Oleoyl-L-Lysine** | 116<br>(91 – 147) | 93.6 ± 1.9 | 269*<br>(180 – 406) | 91.5 ± 1.5 |
| **Oleoyl-L-Carnitine** | 520<sup>a</sup> | 55.5 ± 5.2<sup>b</sup> | 800<sup>a</sup> | 60.5 ± 5.3<sup>b</sup> |
| **Oleoyl-L-Leucine** | 262<br>(195 – 352) | 69.4 ± 3.8 | 619***<br>(464 – 834) | 64.9 ± 3.8 |
| **Oleoyl-L-Tryptophan** | 103<br>(75 – 142) | 98.6 ± 0.3 | 130<br>(97 – 175) | 78 ± 4 |

<sup>†</sup> Activity of bioactive lipids was tested by examining the inhibition of currents produced by application of the glycine EC<sub>50</sub> to WT GlyT2 expressed in *Xenopus laevis* oocytes. Cholesterol depletion was performed by incubating oocytes in 15 mM MβCD for 30 minutes at 32 °C. Values are presented as mean (95% confidence interval) or mean ± SEM (n ≥ 5). Differences between IC<sub>50</sub> and inhibition at 3 μM values between control and treated cells were determined via two-tailed unpaired t-tests. Statistical significance is presented as \* p ≤ 0.05, \*\* p ≤ 0.01, \*\*\* p ≤ 0.001 and \*\*\*\* p ≤ 0.0001.
<sup>a</sup> 95% confidence intervals were unable to be reliably fit.
<sup>b</sup> Values are representative of inhibition at 1 μM due to the chaotropic effects observed with exposure to 3 μM Oleoyl-L-Carnitine.

**Table EV8.**
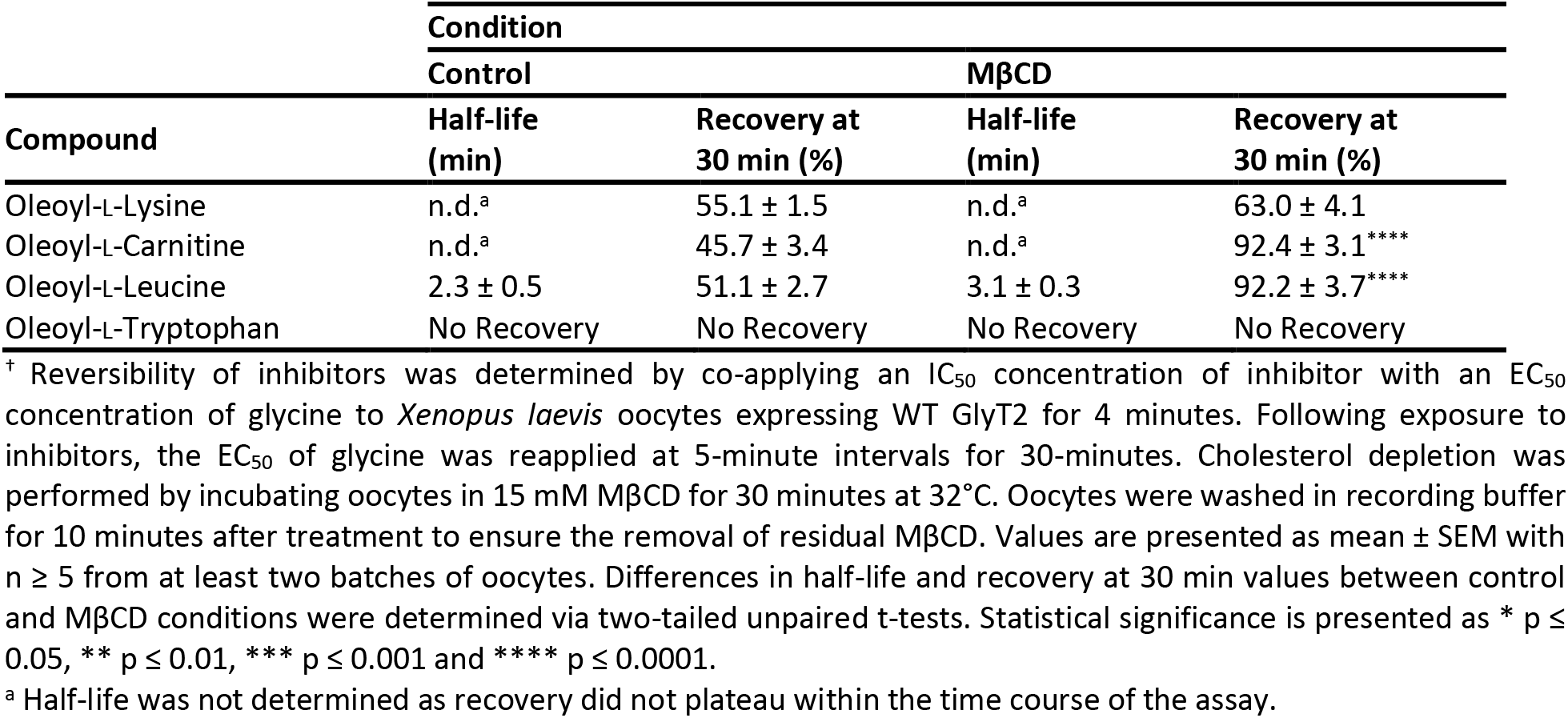
Reversibility of bioactive lipid inhibition of WT GlyT2 expressed in *Xenopus laevis* oocytes pre and post cholesterol depletion^†^.

**Table EV9.** EC_50_ values for glycine transport of WT GlyT2 and CHOL1 site mutant transporters expressed in *Xenopus laevis* oocytes^†^.

| Region | Mutant | Glycine $EC_{50}$ (μM) |
| --- | --- | --- |
| WT | - | 18 |
|  |  | (16 – 21) |
| TM1 | L198A | 28** |
|  |  | (25 – 31) |
| TM5 | Y430L | 19 |
|  |  | (17 – 22) |
|  | Y430F | 12 |
|  |  | (10 – 13) |
| TM7 | T512A | 90**** |
|  |  | (80.76 – 99.83) |
|  | F515V | 7.8* |
|  |  | (6.7 – 9.1) |
|  | F515W | 7.6** |
|  |  | (6.5 – 8.8) |

<sup>†</sup> Glycine $EC_{50}$ values were determined by measuring glycine-dependent currents following application of increasing concentrations of glycine (1 μM – 3 mM) to *Xenopus laevis* oocytes expressing WT and mutant GlyT2 transporters. Currents were fit to the modified Michaelis-Menten equation, and raw currents were normalised to the $V_{max}$ for each cell. Data is presented as mean (95% confidence interval) with $n \geq 3$ from at least two batches of oocytes. A one-way ANOVA determined any differences in mutant glycine $EC_{50}$ values compared to WT. Statistical significance is presented as \* $p \leq 0.05$ , \*\* $p \leq 0.01$ , \*\*\* $p \leq 0.001$ and \*\*\*\* $p \leq 0.0001$ .

**Table EV10.** Inhibitory activity of bioactive lipids at WT and CHOL1 mutant GlyT2 transporters expressed in *Xenopus laevis oocytes*^†^.

| Mutation | Compound | IC<sub>50</sub> (nM) | Inhibition at 3 μM (%) |
| --- | --- | --- | --- |
| Wild-Type | Oleoyl-L-Lysine | 215<br>(136 – 340) | 86.6 ± 1.9 |
|  | Oleoyl-L-Carnitine | 520<sup>a</sup> | 55.5 ± 5.2<sup>b</sup> |
|  | Oleoyl-L-Leucine | 262<br>(195 – 352) | 69.4 ± 3.8 |
|  | Oleoyl-L-Tryptophan | 103<br>(75 – 142) | 76.2 ± 2.3 |
| F515W | Oleoyl-L-Lysine | 689<br>(299 – 1.78 μM) | 85.3 ± 6.4 |
|  | Oleoyl-L-Carnitine | 6.2 μM<sup>a</sup> | 57.6 ± 2.3<sup>b</sup> |
|  | Oleoyl-L-Leucine | 400<br>(307 – 525) | 86.7 ± 1.7<sup>**</sup> |
|  | Oleoyl-L-Tryptophan | 128<br>(64 – 253) | 82.0 ± 3.9 |
| Y430L | Oleoyl-L-Lysine | 863<sup>*</sup><br>(559 – 1.39 μM) | 69.7 ± 4.7<sup>*</sup> |
|  | Oleoyl-L-Carnitine | 1.1 μM<sup>a</sup> | 49.6 ± 2.9<sup>b</sup> |
|  | Oleoyl-L-Leucine | 375<br>(223 – 642) | 63.8 ± 2.6 |
|  | Oleoyl-L-Tryptophan | 57<br>(39 – 85) | 83.5 ± 3.2 |
| Y430F | Oleoyl-L-Lysine | 123<br>(92 – 163) | 92.9 ± 1.7 |
|  | Oleoyl-L-Carnitine | 190<sup>a</sup> | 71.0 ± 3.3<sup>*b</sup> |
|  | Oleoyl-L-Leucine | 203<br>(129 – 316) | 76.7 ± 2.8 |
|  | Oleoyl-L-Tryptophan | 125<br>(91 – 171) | 84.5 ± 3.6 |
| T512A | Oleoyl-L-Lysine | 679<br>(413 – 1.16 μM) | 68.0 ± 1.7<sup>**</sup> |
|  | Oleoyl-L-Carnitine | 1.7 μM<sup>a</sup> | 44.1 ± 2.8<sup>b</sup> |
|  | C18-L-L-leucine | 549<br>(300 – 1 μM) | 50.7 ± 5.7<sup>**</sup> |
|  | Oleoyl-L-Tryptophan | 85<br>(62 – 116) | 74.4 ± 4.5 |

<sup>†</sup> Inhibitory activity of bioactive lipids and ORG25543 was tested by examining the inhibition of glycine currents induced by application of the glycine EC<sub>50</sub> to WT and mutant GlyT2 expressed in *Xenopus laevis* oocytes. Values are presented as mean (95% confidence interval) or mean ± SEM with n ≥ 5 from at least two batches of oocytes. Differences in IC<sub>50</sub> and level of inhibition between WT and mutant GlyT2 transporters were determined by a one-way ANOVA with a Tukey's posthoc test. Statistical significance is presented as \* p ≤ 0.05, \*\* p ≤ 0.01, \*\*\* p ≤ 0.001 and \*\*\*\* p ≤ 0.0001.
<sup>a</sup> 95% confidence intervals were unable to be reliably fit.
<sup>b</sup> Values are representative of inhibition at 1 μM due to the chaotropic effects observed with exposure to 3 μM Oleoyl-L-Carnitine.

**Table EV11.**
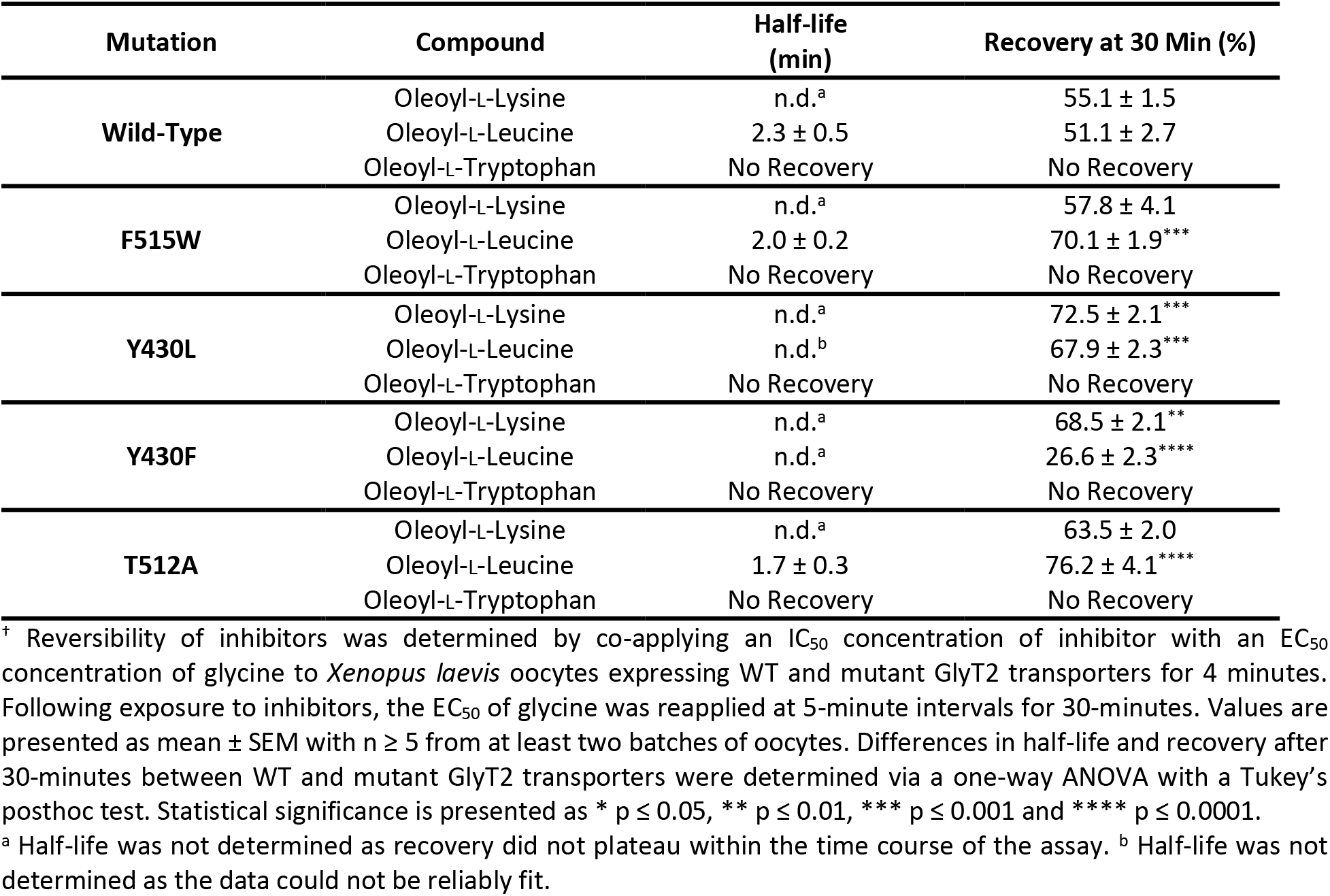
Reversibility of bioactive lipid inhibition of WT and CHOL1 mutant GlyT2 transporters expressed in *Xenopus laevis* oocytes^†^.

| Mutation | Compound | Half-life (min) | Recovery at 30 Min (%) |
| --- | --- | --- | --- |
| **Wild-Type** | Oleoyl-L-Lysine | n.d.<sup>a</sup> | 55.1 ± 1.5 |
|  | Oleoyl-L-Leucine | 2.3 ± 0.5 | 51.1 ± 2.7 |
|  | Oleoyl-L-Tryptophan | No Recovery | No Recovery |
| **F515W** | Oleoyl-L-Lysine | n.d.<sup>a</sup> | 57.8 ± 4.1 |
|  | Oleoyl-L-Leucine | 2.0 ± 0.2 | 70.1 ± 1.9*** |
|  | Oleoyl-L-Tryptophan | No Recovery | No Recovery |
| **Y430L** | Oleoyl-L-Lysine | n.d.<sup>a</sup> | 72.5 ± 2.1*** |
|  | Oleoyl-L-Leucine | n.d.<sup>b</sup> | 67.9 ± 2.3*** |
|  | Oleoyl-L-Tryptophan | No Recovery | No Recovery |
| **Y430F** | Oleoyl-L-Lysine | n.d.<sup>a</sup> | 68.5 ± 2.1** |
|  | Oleoyl-L-Leucine | n.d.<sup>a</sup> | 26.6 ± 2.3**** |
|  | Oleoyl-L-Tryptophan | No Recovery | No Recovery |
| **T512A** | Oleoyl-L-Lysine | n.d.<sup>a</sup> | 63.5 ± 2.0 |
|  | Oleoyl-L-Leucine | 1.7 ± 0.3 | 76.2 ± 4.1**** |
|  | Oleoyl-L-Tryptophan | No Recovery | No Recovery |

<sup>†</sup> Reversibility of inhibitors was determined by co-applying an IC<sub>50</sub> concentration of inhibitor with an EC<sub>50</sub> concentration of glycine to *Xenopus laevis* oocytes expressing WT and mutant GlyT2 transporters for 4 minutes. Following exposure to inhibitors, the EC<sub>50</sub> of glycine was reapplied at 5-minute intervals for 30-minutes. Values are presented as mean ± SEM with n ≥ 5 from at least two batches of oocytes. Differences in half-life and recovery after 30-minutes between WT and mutant GlyT2 transporters were determined via a one-way ANOVA with a Tukey's posthoc test. Statistical significance is presented as \* p ≤ 0.05, \*\* p ≤ 0.01, \*\*\* p ≤ 0.001 and \*\*\*\* p ≤ 0.0001.
<sup>a</sup> Half-life was not determined as recovery did not plateau within the time course of the assay. <sup>b</sup> Half-life was not determined as the data could not be reliably fit.

**Table EV12.** Percentage of the total simulation time in which residues are in contact with Oleyol-L-Carnitine bound in the LAS in atomistic simulations of GlyT2 mutants. Only interactions that occur for >15% of the total simulation time are reported.^a^

| Region | Residue | WT | T512A | Y430F | Y430L | F515W |
| --- | --- | --- | --- | --- | --- | --- |
| TM1 | V214 | 48.6 | 56.1 | 42.7 | 57.6 | 73.6 |
| TM1 | W215 | - | - | - | - | 33.7 |
| TM1 | F217 | 54.8 | 69.0 | 47.3 | 52.1 | 64.5 |
| TM1 | P218 | 62.3 | 46.2 | 68.3 | 67.6 | 60.5 |
| TM5 | Y430 | 44.9 | 19.2 | - | - | 17.3 |
| TM5 | L433 | 24.2 | 35.8 | 33.6 | 45.7 | 16.6 |
| TM5 | V434 | 17.8 | - | - | - | - |
| TM5 | L437 | 53.7 | 60.5 | 55.3 | 25.8 | 34.4 |
| TM5 | R439 | - | - | 89.2 | 82.7 | 81.4 |
| TM5 | G440 | 51.6 | 41.3 | 88.0 | 76.1 | 90.8 |
| TM5 | V441 | - | 15.5 | 44.7 | 18.8 | 25.4 |
| TM5 | T442 | - | 20.2 | 79.8 | 47.9 | 58.3 |
| TM7 | T512 | 29.3 | 31.6 | 29.8 | 24.2 | - |
| TM7 | F515 | 58.3 | 22.8 | - | 39.0 | 22.4 |
| TM7 | A516 | 51.1 | 56.6 | 52.5 | 60.7 | 42.3 |
| TM7 | V519 | 28.2 | 45.9 | 46.4 | 42.3 | 27.9 |
| TM7 | I520 | 60.3 | 69.8 | 75.6 | 77.0 | 72.7 |
| TM7 | V523 | 45.1 | 59.7 | 72.8 | 61.1 | 75.7 |
| TM7 | F526 | 39.4 | 33.7 | 57.6 | 41.1 | 55.5 |
| TM7 | M527 | 53.7 | 59.8 | 40.6 | 46.9 | 58.5 |
| EL4 | V533 | 75.4 | 51.7 | 40.1 | 38.4 | 56.1 |
| EL4 | I535 | 52.4 | 68.4 | 24.6 | 35.2 | 16.1 |
| EL4 | N537 | 35.9 | 75.9 | - | 37.0 | - |
| EL4 | V538 | 38.2 | 32.9 | - | 23.7 | - |
| EL4 | L559 | - | 18.3 | - | - | - |
| EL4 | S560 | 64.1 | 63.5 | 22.6 | 62.4 | 75.6 |
| TM8 | F562 | 97.4 | 93.7 | 98.9 | 99.2 | 98.8 |
| TM8 | W563 | 88.3 | 90.2 | 96.4 | 96.7 | 98.4 |
| TM8 | I566 | 68.4 | 71.1 | 70.7 | 67.1 | 67.6 |
| TM8 | F567 | 75.6 | 55.5 | 65.9 | 64.5 | 88.8 |
| TM8 | M570 | 74.9 | 58.0 | 71.9 | 51.7 | 78.7 |
| TM8 | L571 | - | - | - | - | 27.3 |
| TM8 | T573 | - | - | 33.5 | 33.2 | - |
| TM10 | F629 | 21.4 | - | - | - | 43.5 |

<sup>a</sup>An interaction is defined as a minimum distance between heavy atoms in the residues to be < 4 Å.

**Figure 12 Supplement Table 2.** Percentage of the total simulation time in which residues are in contact with the cholesterol molecule bound near the bottom of the LAS of GlyT2 while Oleyol-L-Carnitine is bound in the extracellular allosteric pocket. Only interactions that occur for >30% of the total simulation time are reported.^a^

| Region | Residue | WT | T512A | Y430F | Y430L | F515W |
| --- | --- | --- | --- | --- | --- | --- |
| TM1 | V205 | 60.7 | 69.5 | 94.7 | 84.6 | 62.7 |
| TM1 | G206 | 35.9 | - | 80.9 | 82.7 | - |
| TM1 | Y207 | - | - | - | 65.3 | - |
| TM1 | A208 | 47.3 | 66.1 | 70.3 | 53.9 | 26.6 |
| TM5 | Y430 | 70.4 | 98.5 | 98.2 | 63.4 | 71.4 |
| TM5 | L433 | - | - | 50.1 | 60.5 | - |
| TM5 | V434 | 44.7 | 62.7 | 61.1 | 53.5 | 57.3 |
| TM5 | L437 | - | 32.6 | 38.9 | 53.9 | - |
| TM7 | T508 |  | 56.0 | 39.6 | - | - |
| TM7 | A511 | 43.8 | 49.7 | 63.5 | 37.6 | 36.0 |
| TM7 | T512 | 47.5 | 50.8 | 77.2 | 60.5 | 58.5 |
| TM7 | F515 | 66.1 | 88.0 | 93.1 | 94.5 | 86.6 |
| TM8 | T573 | - | - | 62.7 | 54.8 | - |

<sup>a</sup>An interaction is defined as a minimum distance between heavy atoms in the residues to be < 4 Å.

## Notes

### Competing Interest Statement

The authors have declared no competing interest.

